# Phage libraries screening on P53 : yield improvement by zinc and a new parasites-integrating analysis and rationale

**DOI:** 10.1101/2023.09.01.555989

**Authors:** Sihem Ben Abid, Emna Ketata, Ines Yacoubi-Hadj Amor, Salma Abdelmoula-Souissi, Lamia Djemal, Aida Koubaa, Raja Mokdad-Gargouri, Ali Gargouri

## Abstract

P53 is a transcription factor that controls a variety of genes, primarily involved in cell cycle and other processes related to cell survival and death. We have isolated peptides targeting P53 (protein and domains) using the “phage display” technique. Interestingly, adding ZnCl2 at 5-10 mM in panning solutions helped to recover more plaque-forming units at least at round one of the screening. Subtractive docking analyses were designed by using a pool of common redundant peptides known as *parasites*. This rationale helped us differentiate between possibly specific and non-specific bindings. We found notable differences in docking characteristics between different sets of peptides either related to different targets or related to zinc-conditions. The set of zinc-related peptides shows advantageous docking profiles: sharper binding for some positions and distinct exclusive bound residues, including the relevant R248 and R273. Zinc would have modulating/helping role in the targeting of protein P53 by phage displayed peptides in addition to an enhancement action on bacterial infection.

## 1. Introduction

The P53 tumor suppressor protein is involved in the cell cycle control and the DNA repair mainly as a transcriptional factor targeting genes involved in these processes [1]. In some cancers, mutations affecting p53 alter several of its activities [2,3]. In addition, P53 is implicated in neurodegenerative diseases; it interacts with tau, specifically the oligomeric tau, and forms oligomers and fibrils in human Alzheimer’s disease (AD) brain [4]. P53 operates at the nuclear and the cytoplasmic levels in several processes and in particular in various cell death pathways such as apoptosis [5] and autophagy [6]. All this makes P53 a major target for research and pharmacotherapies. Peptides are one of those therapeutic molecules that have been tested on P53 with varying degrees of success. The Phage display represents a reliable source of such peptides. After a first basic essay made by Daniels and Lane [7], Tal et al developed a higher throughput phage screening on the p53 protein, by sequencing a great number of peptides [8]. Some of them showed potency on tumor regression in mice. The same research team developed some lead peptides among numerous isolated sequences towards clinical mutant-p53 reactivating application [9]. We also made a contribution consisting in the selection by phage display and study of a heptapeptide called NG7 [10].

Targeting approaches can vary from a direct effect on protein structure-function relation to an interfering binding with partner(s). For wt-p53, researchers have commonly sought to enhance the activity of the protein by dissociating it from negative regulators, thus leading to the accumulation of the functional protein [11,12]. These negative regulators are mainly the E3 ubiquitin ligases MDM2 and MDMX which are causing the degradation of P53 by the proteasome [13]. This applies for example to the PNC-27 anticancer peptide family [14]. Phage display has been a valuable tool in the targeting of this particular axis P53/MDM2-MDMX. Inhibiting or diminishing the protein aggregation is another approach which can concern both wt and mutant p53 [15,16]. Researchers used peptides based on aggregation prone sequences in the targeting of the tumor suppressor p53 protein [17]. The ReACp53 designed peptide caps an exposed sequence prone to aggregation on the protein, inhibits mutant p53 aggregation and restores it [18]. For the mutant forms, usually approaches aim to stabilize the protein. This is the case for the CDB3 peptide derived from the p53 partner 53BP2 which stabilizes the core domain of the mutant protein [19]. Researches also have sought to prevent the oligomerization/tetramerization which can lead to hetero-oligomers composed of wt and mutant monomers.

Another structure-function feature widely used in P53-targeting strategies is the zinc binding property. We recall here that a zinc molecule is maintained by four bonds with four residues of p53 protein which are important for its binding with DNA [20]. Strategies vary from supplementing non functional P53 proteins with Zn ions to using metallochaperone proteins in regulating zinc concentrations [21, 22]. However, to our knowledge no existing results about peptides targeting P53 with relation or depending on the zinc presence or state.

Furthermore, and though the technique of phage display screening has brought real advancement to the targeting strategies and provided interesting ligands, it has nevertheless yielded in some cases non specific interactions. One special category of them often retrieved by separate works was characterized and known as *parasite* phages [23, 24].

We sought in this study to select small peptides having an affinity with human P53 using phage display libraries screening. The ultimate goal is to find new peptide-ligands which could modulate some of the functions of the protein in further future tests. To analyze the results of molecular docking of isolated peptides with protein P53, a rationale of subtractive binding is conceived and followed taking advantage of an obtained pool of redundant *parasite* phage clones. Above all, interesting effects of zinc supplementation during screening and titration are highlighted.

## 2. Materials and Methods

### 2.1. Strains, Vectors, Primers

- ER2738 (New England Biolabs) *E. coli: F’proA+ B+ lacIq Δ(lacZ)M15 zzf::Tn10(TetR)/ fhuA2 glnV Δ(lac-proAB) thi-1 Δ(hsdS-mcrB)5.* Used as host in phage display experiments, the supE suppressor allows the read-through of TAG stop codons.
- BL21 (DE3) (Stratagene): *E. coli* B, F^-^, *dcm, ompT, hsdS* (r_B_^-^m ^-^), *gal* (DE3), protease deficient and contains the T7 RNA polymerase gene, under the control of the *lacUV5 promoter*, integrated in the bacterial chromosome. This strain was used for the expression of the GST-p53 fusion protein as for expressing only GST as control.
- vector pGEX-4T-3 (Addgene_79149) to express only GST protein for alternative screening.
- recombinant pGEX-4T-3-P53: The human-p53 cDNA was previously cloned in the *EcoRI* site of pGEX-4T3 (Addgene_79149) under the lac-trp hybrid promoter that is inducible by IPTG, and fused upstream to the GST (Glutathione S-Transferase) [10].
- Primer used for the sequencing of phages clones: -96 gIII sequencing primer: 5’- CCCTCATAGTTAGCGTAACG, set at 1 pmole/µl.

### 2.2. Peptide synthesis

Two peptides spanning two parts of the p53 coding sequence were synthesized by ProteoGenix (Schiltigheim, France): PD74: PPLSQETFSDLWKLLPENNVLSPLPSQAMDDLMLSPDDIEQWFTEDPGPD is a 50 mer covering the acidic N-terminal region (residues 12-61 of p53), with a MW of 5670.3 Da. SR50: SCMGGMNRRPILTIITLEDSSGNLLGRNSFEVRVCACPGRDRRTEEENLR is a 50 mer covering a part of the DNA-binding central domain (residues 241-291), with a MW of 5624.4 Da. ***Rationale for target selection*:** The PD74 peptide correspondent to the N-terminal part of p53 (12–61) contains many relevant motifs such as the important 9aaTAD. These 9 amino acids transactivation domains are thought to have autonomous transactivation activity in some eukaryotes and may also mediate conserved interactions with general transcriptional cofactors [25]. And SR50 originated from the central DNA-binding domain (residues 241–291) containing many important residues and mutation hotspots.

### 2.3. Target preparation for phage libraries screening

#### 2.3.1. P53 Protein purification

Cultures of BL21 *Escherichia coli* bacteria strains expressing separately the GST and the GST- P53 fusion were prepared as following: culture was conducted at 37 °C until an OD_600nm_ of 0.9, and then induced by 0.5 mM of IPTG for 2 hours at 30 °C and at 250 rpm. After lysis with osmotic method, total protein extracts (soluble fraction) are mixed with Glutathione sepharose beads. Beads are then extensively washed to eliminate unbound proteins and particles. Three pools of GST-P53 fixed beads are used for each type of phage libraries; one per round of screening so that at the end three rounds were done. A pool of GST-fixed beads is prepared and used for subtractive panning which aims to eliminate GST-specific phages in the eluate as well as phages recognizing any other contaminating proteins or particles.

#### 2.3.2. Preparation of p53 derived Peptides targets

Peptide PD74 was prepared as a solution of 2 mg/ml in water. This peptide showed some difficulties to suspend, despite the addition of NH_4_OH and sonication. Dilution of peptide was made in buffer NaHCO_3_ 0.1M pH 8.6 to have the final work concentrations of 10-50µg/ml for phage display. Peptide SR50 was similarly prepared in a final concentration of 50µg/ml. For coating, 150 µl of diluted peptide solution were used per well.

### 2.4. Phage Display experiment and post-screening studies

#### 2.4.1. Phage Display outlines

The kit used was purchased from NEB, specifically “Ph.D.™-7 Phage Display Peptide Library Kit | NEB », with a phage library constructed in M13KE phage vector: 2 x 10^13^pfu/ml supplied in TBS with 50% glycerol. All methods used in this work are based on instruction protocols provided by NE-Biolabs. For example, phage titration is performed by one of two methods: bacterial infection on agar plate or estimation based on the optical density of the phage preparation (phage display book [26]). In the second method, the spectroscopic scanning of filamentous phage solution showed an absorption peak at OD_269nm_. Phage particles per ml = (Adjusted A_269_) x (6×10^16^)/(Number of nucleotides in the phage genome=7222b for M13KE). The measured OD_269nm_ is adjusted by subtracting OD_320nm_ (the baseline).

The other procedures (screening by panning procedure, amplification of phage eluate, collection of phages by using the Polyethylene glycol (PEG)/NaCl procedure, extraction of single stranded phage DNA and physical characterization of positive clones) were also carried out according to the NE-Biolabs protocol manual. Here is a view of mains stages :

#### 2.4.2. Phage display Screening

Using the NEB peptide phage display libraries manual’s protocol; we undertook the screening of three different types of libraries: seven-mer linear libraries, constrained (C-C) seven-mer libraries and the twelve-mer displayed peptide phage libraries. After the binding step which lasts between 10 min and 1 h then washing, elution was carried out with Glycine buffer pH2. Eluates are neutralized. Titration is performed at almost every step. Eluate (E) and amplifiate (A) are titrated to appreciate the screening efficiency and the enrichment of specific phages during and between the different rounds. Titration was performed by bacterial infection with serial dilutions and either drops or plating on Petri dishes giving blue plaques which are counted. Alternatively, the titers of phages were estimated via the OD measured by nanodrop according to the method presented in the book «phage display» [26].

*Protein P53:* Screening of Phage libraries was performed, in micro-centrifuge tubes, starting with a pre-clearing step on GST beads. Thereafter, each pre-cleared phage library is panned on GST-p53 beads; follow the wash and elution steps. An additional condition was introduced as an optimization of the standard protocol. It consists on adding zinc chloride (ZnCl 5-10mM) at almost all stages of phage libraries panning: binding (5 mM), and washing (10 mM). This is done for all rounds and all three phage libraries.

*Peptides:* The panning was carried out on immobilized targets: the peptides are fixed separately to the surface of the 96-well plate. Three peptide phage libraries from NEB “New England Biolabs”: 7, 7 C-C and 12 mers were used, as for the whole protein p53. Titers were checked before each use and were around 10^10^ to 10^12^ pfu/ml. In all the screening rounds, the elution was achieved with 0.2M glycine buffer at pH 2.2 supplemented with 0,1 mg BSA/ml or skimmed milk. Titrating the eluates was done by bacteria infection. Another optimization of the phage display protocol was also performed and consisted on modulating the concentration of Tween 20 concentration, that was increased from 0.1% to 0.5% in the third round of panning, with the aim of increasing the affinity of eluted phages by eliminating non-specific phages which would be weakly bound.

Several phages are collected generally from the third round of each screening (E3). They undergo a first step to verify whether they recognize the p53 target, by ELISA test. ELISA-positive phages are then subjected to DNA extraction and sequencing.

### 2.5. ELISA tests

ELISA tests are carried out between (peptide, protein) targets and isolated phages as follows: the wells are coated with the concerned P53 peptide or protein GST-P53 priorly solubilized digested by thrombin and diluted in PBS. Phages are added at approximate titer of 4.23 10^12^. Following incubation, washing, anti-M13 antibody-HRP (Cytiva Cat# 27942101, RRID:AB_2616587) is added and finally revelation is realized with ABTS substrate. The 96-well plate columns were alternatively coated by peptide/protein P53 target (P) or only blocking buffer (C for Control). Each phage is essayed simultaneously on control and protein wells. A percentage of relative signals will be calculated as: [100-C/P*100]; this gives a qualitative idea on the strength of binding between phage particles and the protein target.

### 2.6. DNA sequencing and bioinformatics analysis

The single-stranded DNAs of phages were extracted and their nucleotide sequences were automatically determined, using primer–96 gIII supplied in the kit. Nucleotide sequence analysis and amino acid sequence deduction were performed using the Bio-Edit Package (v.7.0.5) program.

### 2.7. Molecular Docking

The three-dimensional structures of p53 protein were retrieved from the RCSB Protein Data Bank [http://www.rcsb.org/pdb]. The first structure of p53 used in this study was (PDB: 2LY4. Chain B) from the unstructured transactivation domain (Residues 14–60) of the N-terminus of p53 [27]. Another structure (pdb: 3Q01. Chain A) (p53: p53 DNA-binding (Residues 94–291) and oligomerization (Residues 322–356) domain) [28] was prepared by removing water molecules while the zinc cofactors were retained in the p53 model. The structure of the peptides was constructed using the ACD/3D Viewer software.

Peptide and protein interaction patterns were predicted by Autodock Vina [29]. The grid for ligand conformational search calculations was placed with its centre located at the protein level. For the structure (2LY4 chain B), the docking grid size was 60, 40 and 50 grid points, while the grid centers were -9, 0 and 0. And for the structure (3Q01), the size of the docking grid was 60, 60 and 66 grid points, while the grid centers were designated at dimensions 35, 19 and 74. The best conformations with the lowest binding free energy were selected and analyzed using Discovery Studio 2017 R2 Client.

We should notice that a chimeric P53 protein was designed and optimized by [28] with amino acids substitutions in the DBD to reach a stabilized form. Substitutions present in the 3Q01 structure that we used are: C135V, C141V, W146Y, C182S, V203A, R209P, C229Y, H233Y, Y234F, N235K, Y236F, T253V and N268D. They are presented as such in the docking profiles.

## 3. Results

### 3.1. Zinc chloride addition increases the phage titers at the first round

Regarding phage display against the P53 protein, Zinc chloride (ZnCl 5-10mM) was added during different panning steps, knowing that zinc ions are necessary to the p53 protein to maintain a good conformation and therefore a good functionality. **Tab.1** shows the titers obtained (pfu/µl) in round 1, depending on whether or not ZnCl ions were added. As a consequence of this presence of ZnCl, we noticed for this round 1 an increase in the number of phages (plaques) obtained: ten and seven times more for the 7 and 7 C-C respectively, but especially for the 12-mer peptides library (10^3^ times). This improvement concerned the eluates in the first round. This positive effect of zinc treatment was clearer with the amplifiates (A1). The A1 titers are much higher for the “*with zinc”* condition than for the “*without zinc”* condition and this is true for all three libraries. Finally and for the three libraries, the amplification rate (given by A1/E1 ratio) is higher when adding zinc compared to the standard (**Tab.1).** The two other rounds of panning were not titrated; first because it was not aimed to and this zinc effect on phage numbers was observed by chance. Titration is habitually realized with (recommended) serial dilutions which may substitute to repetition.

*The constrained peptide library has low titers at elution and Zinc chloride barely changes it* With regard to 7 C-C library, numbers of phages recovered at this eluate level are too low with or without addition of zinc; that we cannot conclude on a notable effect of ZnCl. Interestingly, after amplification, there was an important increase in phage titer and this increase is more pronounced with zinc addition.(**Tab.1).**

**Table (1):**
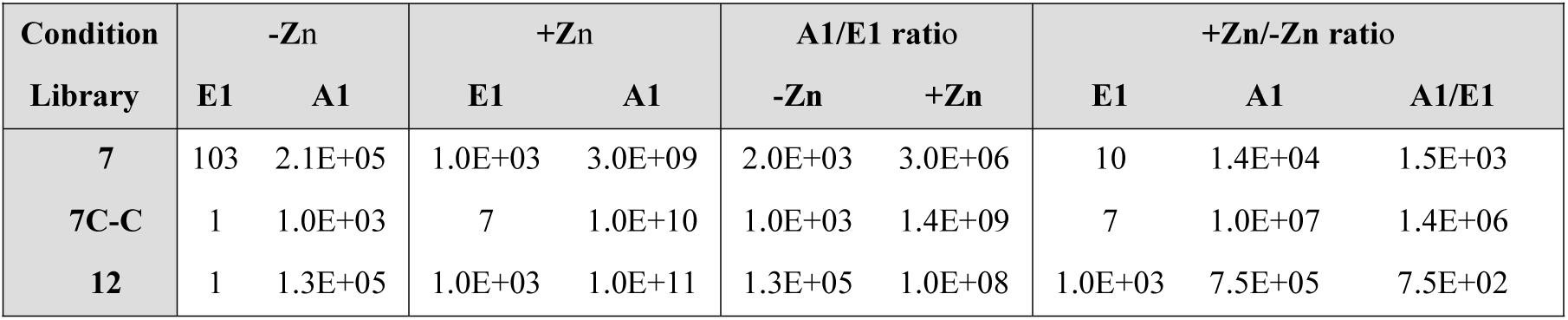
Titles of P53 recognizing Phages. Phage titers (pfu/µl) of the first-round *Eluates* (E) and *Amplifiates* (A) of the three phage libraries. Ratio A1/E1 which reflects the amplification rate is deduced. Libraries are 7 mer (7), 7 constrained mer (7C-C) and 12 mer (12) sized. (Cond: conditions, Lib: libraries).

We think that there were more phages in the eluate of the C-C library than assessed by titration. On the other hand, the low titer of the eluate (E1) of dodecamers’ library, under standard condition, can have different explanations resumed in: technical defects. This will have no repercussion later in the amplifiate titration due to the exponential nature of phage multiplication.

### 3.2. Screening results

#### 3.2.1. New P53-recognizing peptides

Several phages from the third round of screening (E3) were collected and amplified using the bacterial strain ER2738. They underwent a first step to verify whether they recognize the p53 target, by ELISA test **(Fig1. SD)**. ELISA-positive phages are then subjected to DNA extraction and sequencing. Twenty distinct motifs resulting from the screening on the whole GST-P53 protein were obtained: eight patterns (7-mer) and twelve patterns (12-mer) of the correspondent library types (**Tab.2.a-b)**. We mention that sequence (NPNSAQG) that we called 7.3 appears in our previous publication [10]. Phage display against p53-derived domains gives extra motifs: seven 7-mer sequences for the PD74 peptide **(Tab.2c)**, and only two 12-mer’s for the SR50 **(Tab.2d)**.

#### 3.2.2. Screening giving redundant motifs: “Parasite phages”

Other motifs recognizing either PD74 or SR50 or both were separately found, using whether the C-C library or a mixture of the three libraries together: 18 sequences that we called: R0 till R17 **(Tab.2e)**. The remarkable fact was the great similarities between these motifs. However, two consensuses can be sorted out and are shared by the two targets/screenings: NGVEIPP (or GVGIPL) (7-mers) and PFNEPHL (C-C 7-mers).

Some redundant “R” motifs have been already sorted out by our laboratory colleagues in distinct screening works. Researchers, as advanced in introduction, have already reported this type of result “obtaining very frequent repetitive motifs, either in the same screening or in distinct target screenings” [30]. These sequences are known as *Parasites*.

### 3.3. Docking analysis

**Table (2): Phage titles and Lists of P53 recognizing peptides. (a)** Phage titles (pfu/µl) of the first-round Eluates (E) and Amplifiates (A) during screening on GST-P53 protein. Ratio A1/E1 reflects the amplification rate. Libraries are 7 mer (7), 7 constrained mer (7C-C) and 12 mer (12) sized. **(a-b)** Lists of peptides recognizing the P53 protein, of 7-mer and 12-mer sizes respectively. (*) Sequence 7.3: NPNSAQG is reproduced from [9]. **(c-d)** Lists of peptides recognizing p53 domains (PD74 and SR50 respectively) and **(e)** List of redundant motifs (R).

**Table (2):**
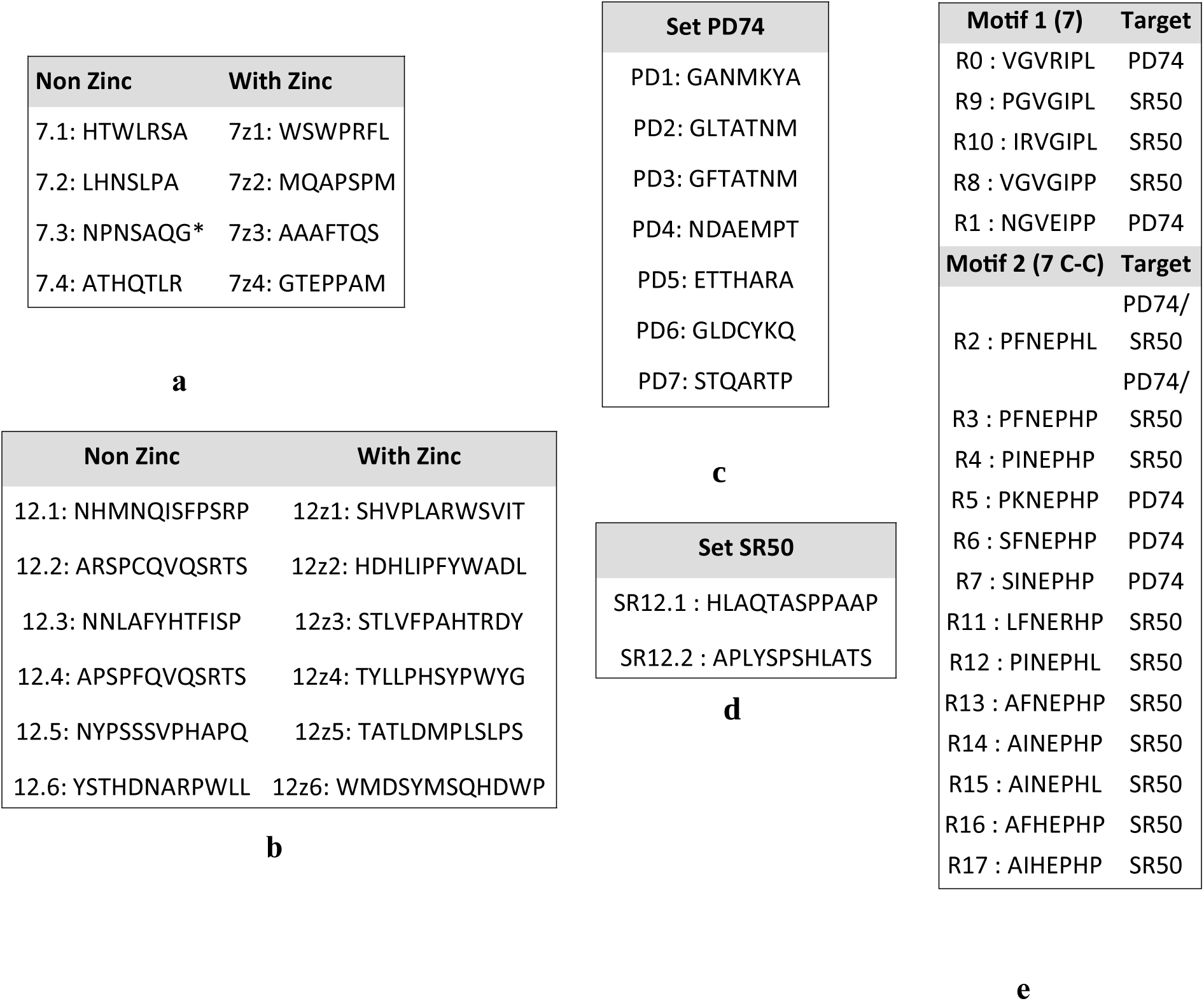
List of phage display obtained peptides. **(a-b)** Lists of peptides recognizing the P53 protein, of 7-mer and 12-mer sizes respectively. (*) Sequence 7.2 NPNSAQG is reproduced from [9]. **(c-d)** Lists of peptides recognizing p53 domains (PD74 and SR50 respectively) and **(e)** List of redundant motifs (R).

**Table 3:**
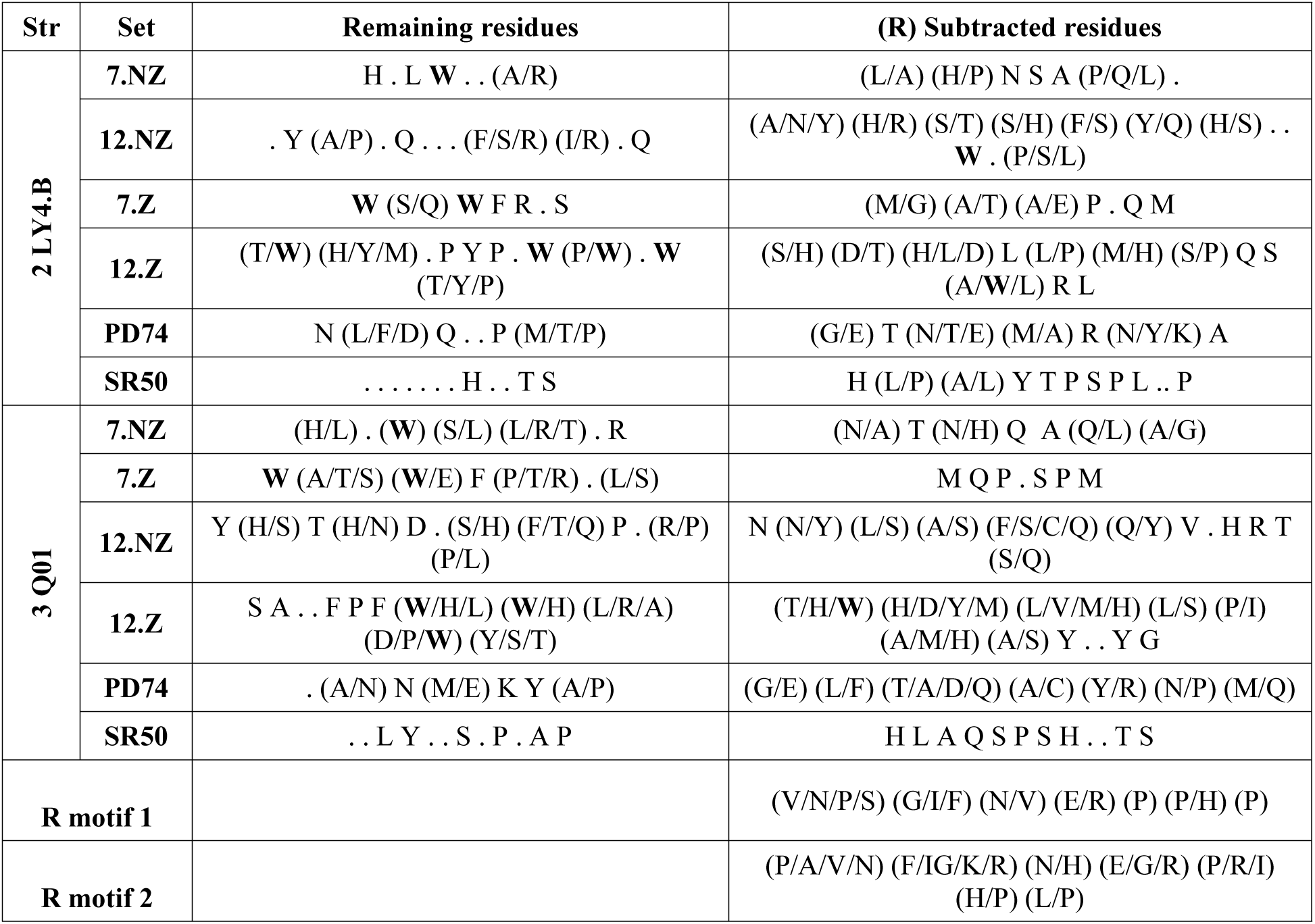
Retained residues Versus subtracted following R correction. (Str : structure.)

A first parameter of docking considered a result is the energy. Values are resumed for all peptides in **(Tab1.SD)**. And we can already remark the superiority of redundant “R” sequences on the others.

#### 3.3.1. Binding profiles in Docking to 2LY4.B

Binding of peptides to 2LY4.B structure has an overall profile of two regions separated by a small area of non-binding in the center A39-M40. The binding frequency ranges from low to high from the ends toward the vertices of these two regions. This is almost the same profile for R set of redundant peptides. **Fig1** shows a comparison once between the 7 and the 12-mer sets (size) and another between the *non zinc* and the *with zinc* states (zinc status) and this is done for the initial form (including the redundant residues) and by subtracting them.

*The ZnCl effect on docking profiles:* Considering the 7-mer sets, the « *non zinc »* profile is closer to that of the *R* set than to that « *with zinc* ». For example, two high contact points S33 and L43 shared between the *non zinc* and *R* sets are weakly present, in the *7 with zinc*. **(Fig.1a)** ZnCl treatment in the 7-mer phage screening made both L43 and P27 buried. It also made the F54 residue newly exposed or available while masking the D57_P60 end reached when docking standard peptides.

For the 12-mer group, the main effect of the ZnCl treatment was to *promote contact with aliphatic/hydrophobic residues* like L45 made accessible as with L35 and V31 or even W23, upstream on the sequence. At the C-terminal end, the zinc treatment, again, *enhances the neutral/hydrophobic residues* (Q52 and W53) and *disadvantages the acidic ones* (E51 and D57). **(Fig.1c)**

In conclusion, *less charges and more aromatics with the ZnCl condition but that was before the R correction.* Moreover, we can observe how the zinc condition refines the docking interactions, especially for the first part of the structure, and makes the contacts sharper and more punctual, also adding more binding-sites.

Redundant bonds frequencies vary from low ∼17% (region 46-56) to high (33, 50 and 67%) in region (29-45).

If we correct by subtracting the common residues from the set *R*, there remain fewer contacts; they flank regions of strong redundant binding or alternate less frequent but ubiquitous bindings. In terms of distribution, differences can be observed between all sets. While the 7*-mer non zinc* positions are spread out (from 27 to 61), the *7-mer with zinc* has only four close contacts at S37, D41, L45 and D45, thus a localized binding. We also observe that “*12 non zinc”* which covers almost the same region than “*7 non zinc”* (middle and C terminal), has a shift compared to “*12 with zinc”* which prefers to bind to the N terminal of the structure. We cite S20, W23, P27 and V31 and far downstream, W53. **(Fig.1h-j)**

#### 3.3.2. Binding profiles in Docking to 3Q01

##### Some relevant residues are exclusive to the “plus zinc” set

**Fig 2** shows the docking profiles of all peptide sets whether including the R redundant residues or after subtracting this redundancy from interactions. On the DNA Binding Domain (DBD), the interaction of all sets recognizing P53 protein can be broadly summarized in three regions: 96-165, 200-233 and 248-291. Considering the docking frequencies, an ascending order is observed: “*12 non zinc”* (16,67%) then both 7-mer sets at 25% but the “*7 with zinc”* has higher binding at some points. And finally, the “*12 with zinc”* set binds at 20%, 33% and 50%. **(Fig 2)**

After *R correction*, only few contacts are lost for “*12 with zinc”* and these are the least frequent binders. The “*12 non zinc”* set loses many binding points. For the 7-mer’s, it is the “*with zinc”* set which loses the relevant binding points which are the most frequent.

When comparing the docking profiles of the 7-mer sets with the two 12-mer sets, differences are obvious **(Fig.2.a-d)**. Note that the former only bind with interval (97-267). The 12-mer sets bind everywhere but seem to favor, in particular the “*12 with zinc”,* the region (248-291) in the extreme DBD.

The 7-mer sets of « *Non Zinc* » and « *With Zinc* » nearly coincide in docking positions except for a few offsets or few distinct contacts. The difference between the two zinc-conditions is clearer for the 12-mer size.

Fewer binding positions are observed for “*12 With Zinc”* especially in the DBD. However, these docking points are stronger than those of “*12 Non Zinc”*: with higher frequencies and reinforced by doublet’s interactions. **(Fig.2.a-d)**

For the « *Non Zinc* » pool, 7-mer and 12-mer almost coincide in bound sub-regions. For the «*With Zinc* » pool, the shift of the 12-mer set to the right (the C-terminal part) compared to the 7-mer is evident.

For SR50 and after subtracting *R*, the docking concerns only a few contacts that all except E271 are probably non-specific since outside the SR50 location. We noticed that the SR50 set shares almost all of its few positions with *12 with zinc*. There is possibly a similarity between the conformations of the synthetic SR50 and the same peptide in the « *With Zinc* » P53 protein; zinc would have exposed some regions there. **(Fig.2.g-h)**

The PD74 group (control here) shows low binding frequencies for most involved residues in the DBD. The frequency (∼14%) is a little higher than those of R set. When R-corrected, only a few contact residues remain which still can be considered as non-specific. **(Fig.2.g-h)**

Finally, by clearing all sets from R residues but also SR50 few positions (non specific), we obtain for the total set of peptides targeting the fixed P53 protein, the following positions in decreasing order of frequency: L45 (30%), V31 (25%), P27 and D41 (15%), W23, E28, N30, W53 and P60 (10%), S20, L26, Q52, E56 and G59 (5%). With residues of the *non zinc* pool being: E56, G59 and P60, those from the *with zinc* pool being: S20, W23, L26, N30, Q52 and W53 and those common to both being: E28, P27, V31, D41 and L45. **(Fig.1h)** The *PD74* set now binds 2LY4.B on four contacts : V31, L45, I50 and W53 all present at 14%. Details of docking structures figure in **(Fig.1.1-5.SD)**.

**Figure 1:**
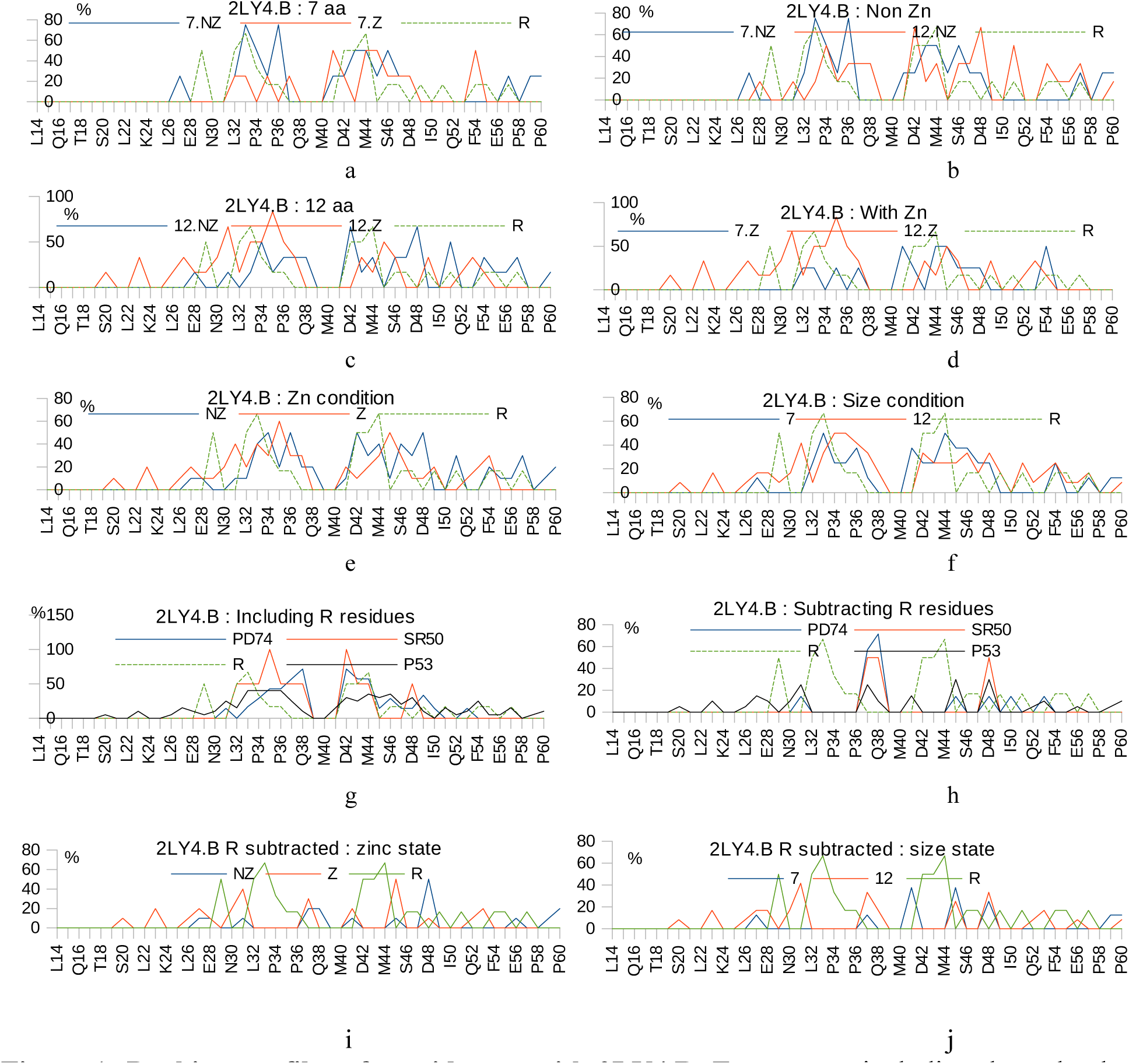
Docking profiles with 2LY4.B. Two ways : including the redundant (R) residues **(a-g)** and with R subtraction **(h and j)**. Comparing the zinc states **(a, c and e)**, comparing the two sizes **(b, d and f)**. And comparing separately the zinc states and the size states on R subtracted condition **(i and j)**.

For the C-terminal region, most of the binding contacts are probably non specific/exclusive: found with *R* set or *PD74* or *SR50*; a single (specific) residue (A353) can be retained for the 12-mer set.

Some residues are rather exclusive to the zinc condition: A129 (7-mer), K132, Q165, R248, P250, E271 and R273 (12-mer). Thus, the relevant residues like the R248 or R273 hotspots are finally reached. We even notice that this binding occurs through the doublets 129/132, 248/250 and 271/273 which means that it is rather strong. R248 is bound by 12Z2 (by W9) which also connects P250 by Y8 and R273 by A10. 12Z3 binds R248 (by L8) and R273 (by S1) which binding is consolidated by E271 bound by D11.

This 12Z3 has an interesting binding involving both the DBD and the C-terminal region; the latter reached by the trio F5-P6-H8. 12Z5 binds R248 via its L3; it also connects to the C terminal region via its L10-P11 pair. Details of docking structures are presented in (Fig.2**.1-14.SD).**

**Figure 2:**
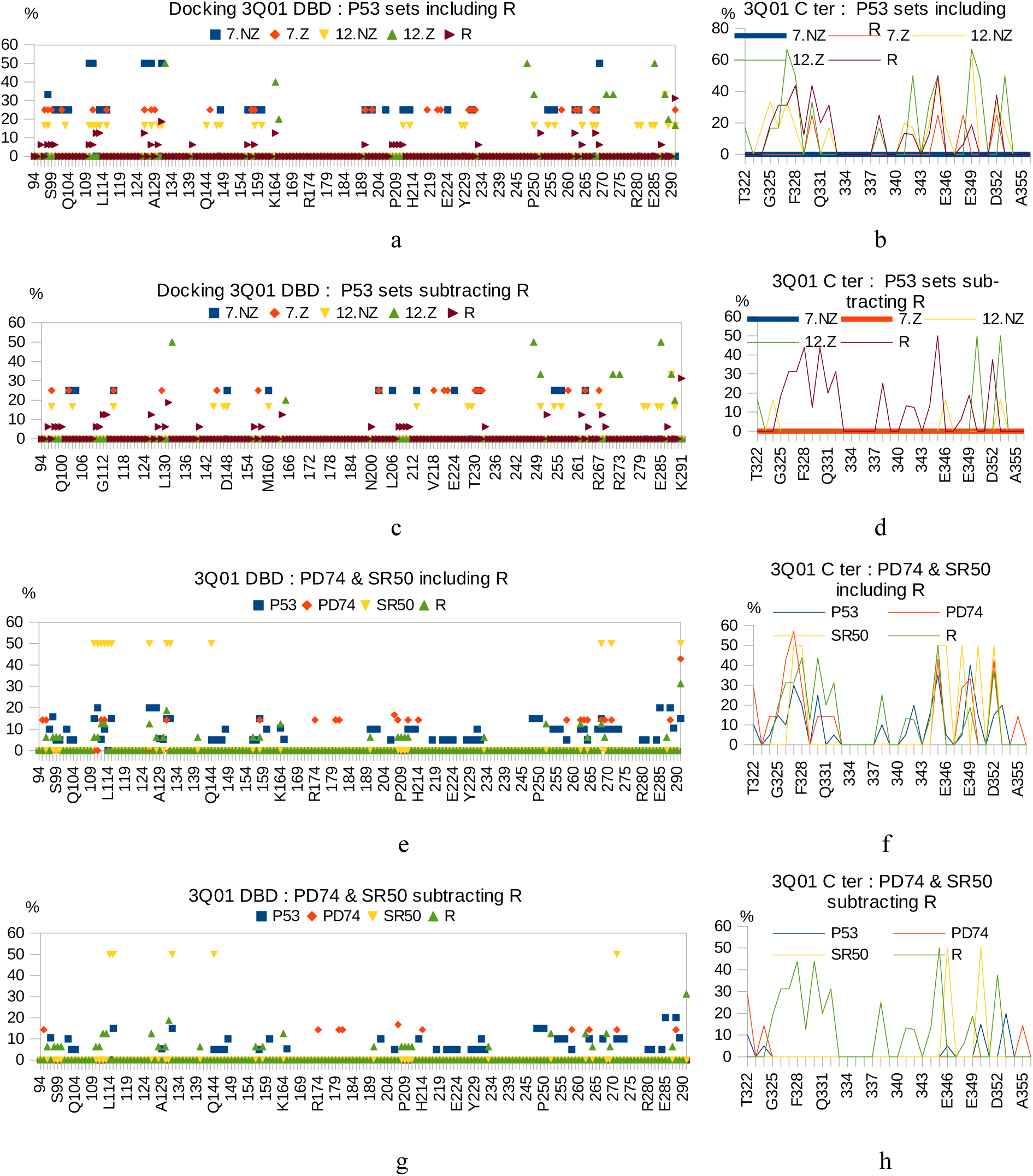
Docking profiles with 3Q01. Profiles related to the DBD (left) and the C ter (right) separately, for : the P53 targeting sets, once including common residues with redondant set R **(a-b)** and once subtracting them **(c-d)**, and for the P53-domains targeting sets, including R residues **(e-f)** and subtracting them **(g-h)**.

### 3.3.3. Test of interaction between peptides and protein P53 by ELISA

Fig.1SD shows a representative ELISA test realized to assess interaction between phages and P53 target. We notice this: 12.6, 12.4, 7Z3 and 7.4 which show a good docking give good response at ELISA. 12Z4 which mainly docks on N-ter shows similarly a good signal. Curiously, some good binders like a series of “*12 with zinc”* show weaker signals. This is not problematic, since “protein-peptide” or “protein-protein” interactions are rather weak in nature [31].

### 3.3.4. Suggestion of retained sequences after R correction

Details of retained positions and subtracted ones for the distinct sets of peptides are summarized in **Tab 3**. We notice the presence of more Trp residues in the retained sequences than in the subtracted positions. These Trp residues overabundance can result from a bias as described in [31]. It has been advanced that this Trp bias in phage display output indicates that the corresponding positions are important to the interaction with peptide ligands [31].

### 3.3.5. Distribution of amino acids classes on target contacts with and without correction

We studied the amino acids classes distribution among the bound residues found by docking (Fig.3). Overall for the distinct regions, we remark the extinction of aromatics with the R set subtraction except with some cases. Thus aromatics seem to characterize the redundant binding. The DBD shows less change in the distribution of amino acids classes than both the N-terminal and C-terminal regions. The R subtraction did not affect the initial distribution. Redundant binding does not have preferential classes of amino acids (except the aromatics) here for the DBD.

**Figure 3:**
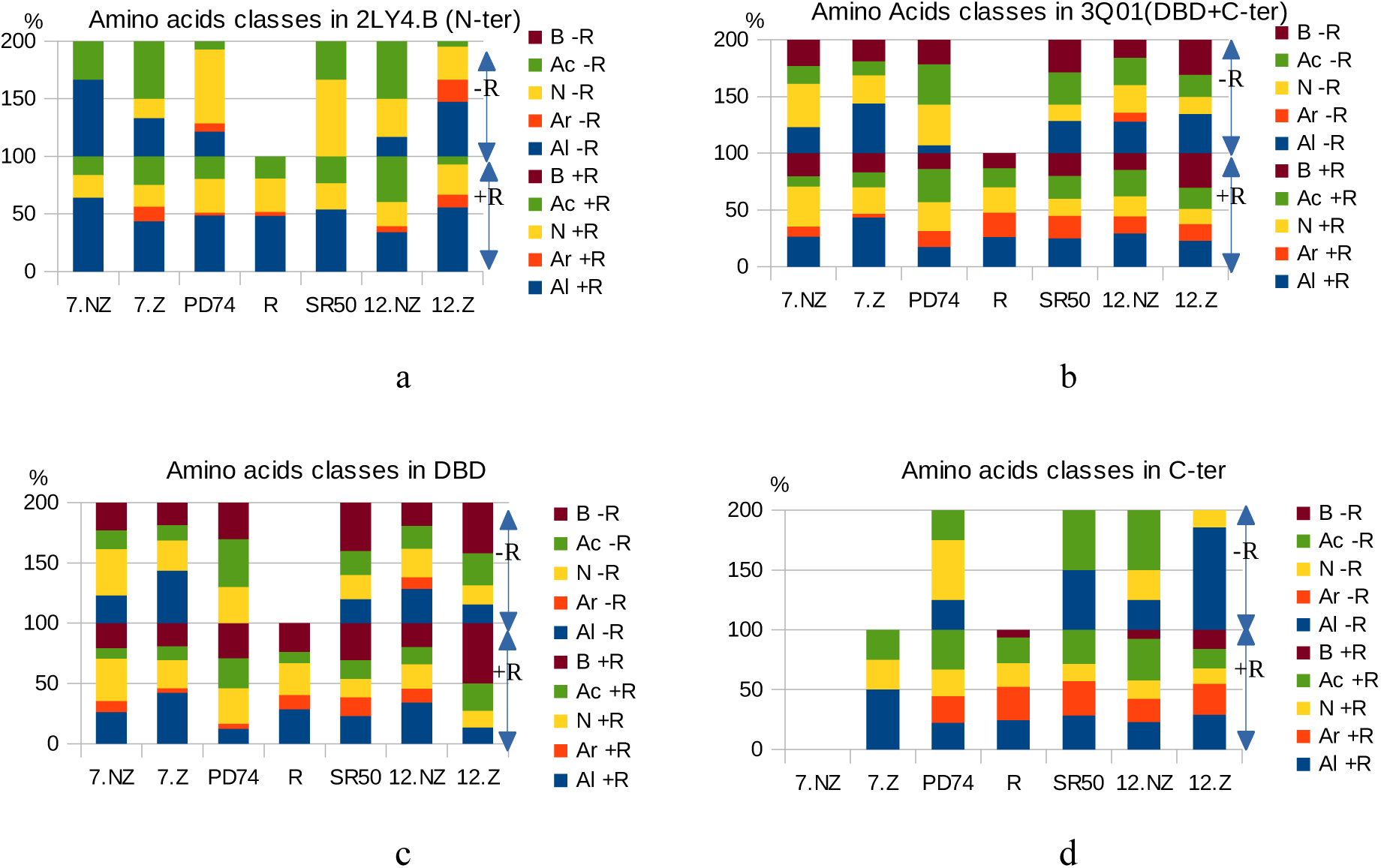
Distribution of amino acids classes on target contacts. Two conditions : including redundant R set positions (+R) and without them (-R). Amino acids classes : B: basic, Ac: acid, N: neutral, Ar: aromatic and Al : aliphatic. At the y axis, % of presence of amino acids classes is cumulative here.

This is not the same for the N-terminal and the C-terminal regions where some classes can disappear with R correction. We notice how the “*12 with zinc”* set binds to basic residues class in the DBD more than any other set (Fig.3**.c)**; this is likely inherited from its higher acidity.

### 3.3.6. Docking analysis of individual peptides after R correction

We examined the docking structures **(see SD)** obtained for all our isolated peptides with a knowledge of their docking profiles (Fig 4) once stripped of redundant common positions (from the *R* set), as well as the control non specific sets *SR50* (2LY4) and *PD74* (3Q01).

**Figure 4:**
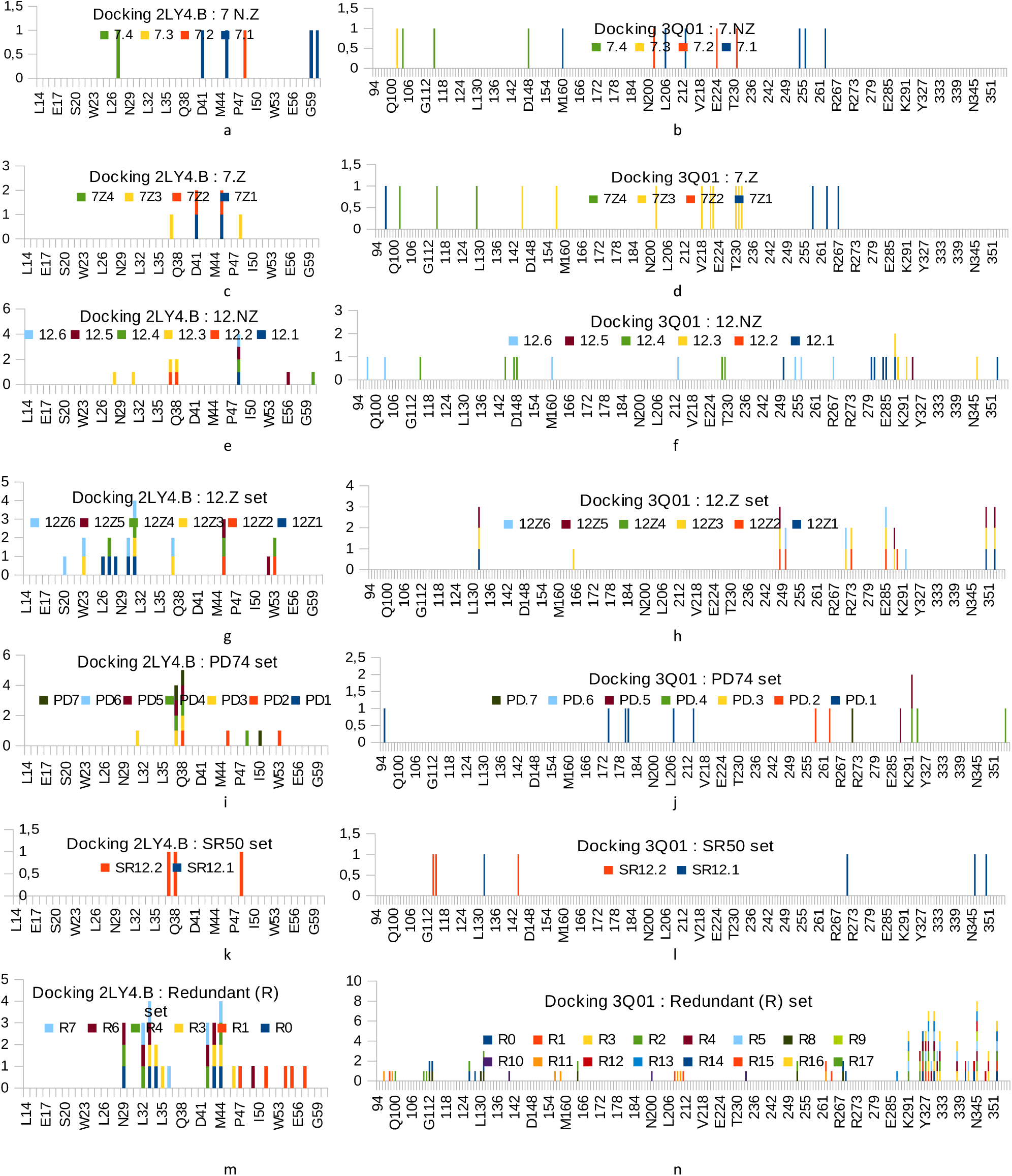
Docking profiles of individual peptides after R correction. Docking points of Redundant peptides are subtracted from the other sets to show only the specific interactions. At left, are presented dockings to 2LY4.B and at right are presented dockings to 3Q01. (a,b) : 7 Non Zinc, (c,d) : 7 With Zinc, (e,f) : 12 Non Zinc, (g,h) : 12 With Zinc, (i,j) : PD74 recognizing set, (k,l) : SR50 recognizing set and (m,n) : Redundant (R) set.

## Individual peptides docking on 2LY4.Chain B

### 7-mer set of P53-recognizing peptides (Fig.1.1.SD)

Peptide 7.1: HTWLRSA shows an interesting docking, making a loop between region (41-47) and the end (57-60). It is the single motif that attaches to this end (57-60). 7Z1: WSWPRFL and 7Z2: MQAPSPM dock at the same place (41-54) and almost the same residues. Structurally, this makes a loop spanning helix (46-51) and connecting the region before it to the C-terminus.

However, a part of the binding (P47/F54) is ignored if we correct by subtracting the R residues. The *remaining bindings are: (D41-H1/W3, L45-H1, G59-R5 and P60-L4/R5) for 7.1, (D41-W3/R5 and L45-W1/S2) for 7Z1 and (D41-Q2 and L45-Q2) for 7Z2*.

### “12 non-zinc” set (Fig1.2.SD)

Peptide 12.3: NNLAFYHTFISP docks in the region (28-38) but two positions remain after the double correction (subtraction of *R* and *SR50*), E28 (N1) and V31 (A4). Additionally, 12.4:APSPFQVQSRTS, 12.5: NYPSSSVPHAPQ and 12.6: YSTHDNARPWLL dock to similar regions and in the same way making a loop spanning the (46-51) helix and connecting the front region to the C-terminus. However, the correction leaves only 12.4 and 12.5 with a P60 (P4) and E56 (Y2) positions respectively.

### “12 with zinc” set (Fig.1.3.SD)

Peptides 12Z6: WMDSYMSQHDWP (20-37), 12Z3: STLVFPAHTRDY (23-43) and 12Z1: SHVPLARWSVIT (26-35) dock with the indicated similar regions in the N-terminal part. The *peptide 12Z6 shows an interesting docking:* a close link in particular *with residues S20 (W11), W23 (Y5, W11), N30 (P12) and V31 (W1, P12) which are retained*, linked by the peptide residues indicated. 12Z1 has a rather delimited (strong) docking region distant from any secondary structure including that at the N-terminal extremity (14-20). The *L26P27E28-N30V31 region is retained after correction. The binding is as follows: L26-P4, P27-P4, E28-W8/S9, N30-T12 and V31-H2. 12Z3 retains two positions: W23 (Y12) and V31 (Y12)*.

Otherwise, 12Z4: TYLLPHSYPWYG (27-35 and44-54), 12Z2: HDHLIPFYWADL (29-36 and 43-53) then 12Z5: TATLDMPLSLPS (34_36 and 45_52) dock with the large regions indicated. These regions are more close to the C-terminus, spanning the second red/green structure (46-51). The 12Z2 and 12Z4 peptides have more spatial and three-dimensional docking than the 12Z5 due to their sequences. They contain Imidazole, Proline and aromatic radicals and residues. However, after correction by *R* residues subtraction, only 12Z4 saves its spread docking with four positions: P27 (T1), V31 (Y2), L45 (P9) and W53 (Y8). This brings out the pattern T1Y2-Y8P9 as relevant. 12Z2 joins 12Z5 by linking only two positions: L45 and Q52(12Z5)/W53(12Z2). The binding is composed of: (L45-P6 and W53-W9) for 12Z2 and (L45-L4 and Q52-T1) for 12Z5.

### PD74 set (Fig.1.4.SD)

Regarding this group of peptides, they all dock with a region in the (32_50). And some of them interact with the helix (46-51): PD1 (GANMKYA), PD2 (GLTATNM), PD4 (NDAEMPT), PD3 (GFTATNM) and PD7 (STQARTP). PD5 (ETTHARA) has a concentrated docking located in a small region (34_44). PD6 (GLDCYKQ) seems to have fewer bonds than the others: 2 liaisons with 2 residues, P36 and L43. The remaining interactions after eliminating redundancy (and non-specificity) are: (L45-L2 and W53-G1/L2) for PD2, (V31-F2) for PD3, (I50-P7) for PD7. Most of the interactions of this set therefore seem non specific almost like the ubiquitous *R* set.

### R and SR50 sets (Fig.1.5.SD)

The binding of the R set (6 representative motifs) remains the same since no corrections have been made. Only R1 (NGVEIPP) has a bond shifted to the right (C-ter end) (P47-E51-F54-T55-D57). The rest of the group docks as follows: (N29-S33-P34-L43-M44) for R0, (S33-P34-L35-L43-M44-S46) for R3, (N29-L32-D42-M44) for R4, (N29-L32-S33-D42-L43-D49) for R6 and (L32-S33-P36-D42-M44) for R7.

This set reaches helix (42-49). Regarding the control docking of *SR50* set, only SR12.2 has a remaining bond itself considered as non-specific: (S37:S7-Q38:S7/H8-D48:H8/T11/S12).

### Analysis of retained docking interactions in 3Q01

Some peptides are simply rejected after R correction, we cite: 7Z2, 12.2, 12Z4 although other interactions can still be rejected since they are found with the PD74 or SR50 sets (outside the 241-291 region); 12.3 and 12.5 can therefore also be ignored.

### Individually for each peptide, as shown by (Fig 4), we have

7.1 retains: M160-H1, L206-L4, R213-W3, I254-H1 and T256-H1. 7.2 retains: R202-S4, E224-L1/Q5, T231-N3. 7.3 retains only T102-S4. 7.4 retains: Q104-R7 and D148-R7. 7Z1 retains: V97-W3 and R267-W3. 7Z3 (AAAFTQS) retains a good number and distribution of interactions: L145-F4, V157-F4, R202-T5, V218-F4, E221-S7, P222-S7, T230-F4, T231-A2 and I232-F4. Obviously the F4 of this peptide is highly interactive; with essentially hydrophobic bonds. 7Z4 retains: T102-E3 and A129-T2.

For 12.4, are retained: V147-S3, D148-S3, D228-F5 and Y229-F5. For 12.1: R280-H2, D281-N1/S7, T284-N1/N4, E285:OE2-S7/F8/P9, P250-P9 and A353-N4. For 12.6: V97-L12, M160-H4, R213-D5, I254-H4, T256-H4 and R267-Y1/S2/T3. 12Z1 retains one position: A353-W8. 12Z2 retains all its interactions: R248-W9, P250-Y8, R273-A10, E285-A10 and L289-F7. 12Z3 retains: Q165-Y12, R248-L3, R273-S1, N285-S1 and A353-H8. 12Z5 retains: R248-L8, A353-P11. 12Z6 retains: Pro250-W11 and E285-W11.

We assemble the retained residues in peptides for all these sets, resulting in: for the 7-mer set: 7.1 (H1W3L4), 7.2 (L1N3S4Q5), 7.3 (S4), 7.4 (R7), 7Z1 (W3), 7Z3 (A2F4T5S7) and 7Z4 (T2E3). For 12-mer motifs: 12.1 (N1H2N4S7F8P9), 12.4 (S3F5), 12.6 (Y1S2T3H4D5), 12Z1 (W8), 12Z2 (F7Y8W9A10), 12Z3 (S1L3H8Y12), 12Z5 (L8P11) and 12Z6 (W11).

## 4. Discussion

We performed p53 targeting by phage libraries. Interestingly, the addition of ZnCl_2_ proved effective in improving the number of pfu obtained for eluates at round 1. Reasons for this phage titers improvement can be variable and need to be known. It has been found that metal cations: iron, magnesium, and calcium enhance the phage killing of bacteria cultured on treated blood medium [32]. That was specific to some phage over others and was true for some bacteria strains over others. Researchers gave a rich discussion about possible explanations for this effect of metal cations (Fe, Mg and Ca) on phage infection and killing of bacteria ; which have been already observed and studied in standard bacterial culture media [32]. In sum, cations can affect the polarization of bacterial membrane, interfere with the DNA entrance into cells or enter into the composition of phage proteins (cofactors) and the intracellular synthesis of phage, etc [32]. These results and explanations can probably apply on Zn cation and we think that Zn would have improved phage infection and/or stability at the titration steps, especially when ZnCl2 is present at eluation at the designed concentrations. This possibly means that eluates’ titers are underestimated at the standard conditions (without Zn) because of lack of infection (or stability) of phages. The presence of Zn would only facilitated the infection of phages already obtained ; not improved the panning (elution) output itself.

For amplifiates, a positive effect of Zn on phage titers can be explained by the same way : Zn enhances infection of bacteria. However, the enhancement is relatively less important than for eluates since ZnCl2 was already diluted first by the passage of low volumes of phage eluates into larger volumes of bacterial cultures during amplification and possible use of the salt by bacterium itself. Then consecutive steps of phage recovery, concentration by PEG, centrifugation, etc finished probably by eliminating any Zn present. So here any positive effect of Zn was probably limited to the amplification phase and did not concern the titration infection which followed it.

The presence of metal ions (Zncl) was not suffisant enough to enhance infection at titration for C-C library eluates. With or without ZnCl in solutions, eluates titers were low while amplifiates were much higher for both conditions. The low eluates titers probably don’t reflect low phage charges. The constrained C-C phages were obtained in a form/state which can not infect bacteria properly following elution. A possible explanation for this can be an effect of low pH elution on Cys residues and disulfide bonds. A reduction of thiol entities of the introduced pair of Cys flanking the exposed peptides may affect the adhesion of phages to bacterium by introducing charge modifications or recreating disulfide bonds intra-pIII. But this changed after amplification and phage recovery by PEG concentration: Cys state may have changed upon different treatment, cell uptake and dilution. And we retrieved expected important titers, yet again with a superiority for the “ZnCl” pool.

At another level, Zn presence would have led to conformational changes that occur on the P53 protein, specifically by exposing more binding sites or residues. The interaction patterns, as well as the binding sites, of the two types of peptides under the « *with* and *non zinc* » conditions were different as modeling studies have confirmed. The most relevant fact is the ability of some *with zinc* peptides to specifically target residues R248 and R273. The presence of zinc must have made these positions accessible. This may provide a piece of explanation to the fact that supplementation of some p53 mutants (R175H and R273H) with zinc recovered their functionality [21] as well as to the zinc metallochaperones capacity to restore the p53 and recover its functions [22].

The important region for MDM2 binding on p53 is known to be (15-29), with the triad (F19, W23 and L26) forming the amphiphilic α-helix in the complex [33]. Peptides 12Z6 and 12Z3 both reaching residue W23, and 12Z1 by binding tripeptide 26-28, are good potential interfering ligands. Numerous data support a crucial role of zinc in “conformational” regulation of P53 [34,35]. The DBD depends on the availability of zinc for its appropriate folding and therefore its function. Its affinity for zinc is described as one of the highest of all time. In the cell, the P53 protein is maintained in a folded and an unfolded states by certain mechanisms regulating zinc [34,35]. Therefor, since our results and this literature, adding zinc in peptide screens and tests can become a systematic choice, as well for P53 and other targets.

In the N-ter structure, we can notice special features such as a symmetry of binding around residue L45, common to *non zinc* and *with zinc* categories however reversed between them. This symmetry engages between others the ubiquitous positions since it disappears when R corrected. Researchers have stipulated a mechanism of interaction between transactivation domain (TAD) of the P53 protein and its partner the replication protein A (RPA), which consists of a conformational transformation of this TAD upon binding. The originally largely disordered TAD in solution, folds via its residues 37–57 into two amphipathic helices, H1 and H2 [36]. Moreover this TAD conformation seems to be modulated by some ligand binding (epigallocatechin gallate EGCG) and cancer-associated mutations N29K/N30D [37]. This ligand binding effects are mainly achieved through nonspecific interactions [37]. Here, it can be said that this domain can have two distinct behaviors in the absence and in the presence of zinc ions. We believe that peptides screening and binding studies can bring some understanding to this structure-function knowledge of the TAD region.

Despite the impressive number of peptides already isolated, our sequences are all new peptides interacting with p53. Meanwhile, we note that several sub-motifs of our sequences have been found approximately in other motifs isolated by other authors [7,8]. We quote patterns as RVVG/VGVR, ALTH/TH or LHA, FSSP/FISP, PFTNN/PFQ, which we find respectively in their results and in ours. This raises the question of the ubiquitous binding already encountered intrinsically.

The knowledge of these parasitic or redundant/ubiquitous motifs and (sub)motifs is very important for studies of any: peptide screening, isolated peptides or ligand-target interactions. Herein from the docking profiling study, we can even make hypothesis which P53 residues are most likely to bind non-specifically to any other molecule(s) or peptide(s) tested.

We used these ubiquitous sequences (named R set) in our study; the characteristics of their interactions led us to choose to subtract this “non-specific” binding from the others, resulting in cleared/corrected versions of all obtained sets of peptides. Retained interactions remain interesting even if they are few in number. Further confirmation with techniques like alanine substitution and biophysical binding studies (RMN…) will help us assess the accuracy of this rationale of redundancy (parasite) subtraction. This is encouraging to further characterize these peptide sequences (original and corrected), their modes of binding to targets, to synthesize them and to test them in cellular and animal models.

In summary, we report an important improvement of screening toward P53, quantitatively and qualitatively with zinc addition in the protocol. This is put in evidence at both the biological interaction level (infection) and the structural study level (docking). Besides, we describe a new approach to refine the screening of peptides potentially applicable to other targets. In conclusion, we have provided new insights into the phage library screening method.

## Author Contributions

S.B.A and A.G conceived the study. S.B.A, I.Y.H.A, S.A, L.D, A.K and A.G conducted the protein expression and purification. S.B.A and A.G conducted the phage display screen. S.B.A, N.K and A.G realized the Nucleic acids sequencing. E.K performed the Docking software manipulation and S.B.A analyzed Docking output. R.M.G, I.Y.H.A, S.A, L.D, A.K and A.G prepared ressources. S.B.A made the first writing-review and editing ; S.B.A and A.G reviewed the manuscipt. A.G supervised and administered the project. A.G acquired funding. All authors have been read and agreed the published version of the manuscript. Note : Authors disclaim any use of *generative artificial intelligence (AI) or human-assisted editing services*

## Informed Consent Statement

Not applicable.

## Data Availability Statement

All data generated or analysed during this study are included in this published article [and its supplementary information files]. The datasets generated and/or analyzed during the current study are available in the [figshare] repository, [<u>10.6084/m9.figshare.24045519</u>]. Besides, the datasets used and/or analyzed during the current study available from the corresponding author on reasonable request.

## Acknowledgments

We thank members of the LBME as well as members of the Unité d’Analyses at the CBS for any technical help. Arnaud Bondon is warmly thanked for having received S. Ben Abid in his laboratory at the University of Rennes I and for his encouragement. *This publication is dedicated to the memories of Basma Hentati and Abdellatif Maalej teachers/researchers at Faculty of Sciences of Sfax*.

## Additional Information

## References

1. Engeland, K. Cell cycle regulation: p53-p21-RB signaling. Cell Death Differ 29, 946–960 (2022). 10.1038/s41418-022-00988-z

2. Baslan, T. et al. Ordered and deterministic cancer genome evolution after p53 loss. Nature. 2022 Aug;608(7924):795-802. doi: 10.1038/s41586-022-05082-5. Epub 2022 Aug 17. PMID: 35978189; PMCID: PMC9402436. https://pubmed.ncbi.nlm.nih.gov/35978189/

3. Kim MP, Zhang Y, Lozano G. Mutant p53: Multiple Mechanisms Define Biologic Activity in Cancer. Front Oncol. 2015 Nov 11;5:249. doi: 10.3389/fonc.2015.00249. PMID: 26618142; PMCID: PMC4641161.

4. Farmer, K.M., Ghag, G., Puangmalai, N. et al. P53 aggregation, interactions with tau, and impaired DNA damage response in Alzheimer’s disease. acta neuropathol commun 8, 132 (2020). 10.1186/s40478-020-01012-6

5. Aubrey BJ, Kelly GL, Janic A, Herold MJ, Strasser A. How does p53 induce apoptosis and how does this relate to p53-mediated tumour suppression? Cell Death Differ. 2018 Jan;25(1):104–113. doi: 10.1038/cdd.2017.169. Epub 2017 Nov 17. PMID: 29149101; PMCID: PMC5729529.

6. Ma S, Kong D, Fu X, Liu L, Liu Y, Xue C, Tian Z, Li L, Liu X. p53-Induced Autophagy Regulates Chemotherapy and Radiotherapy Resistance in Multidrug Resistance Cancer Cells. Dose Response. 2021 Oct 8;19(4):15593258211048046. doi: 10.1177/15593258211048046. PMID: 34646092; PMCID: PMC8504250.

7. Daniels, D.A, Lane, D.P. The characterisation of p53 binding phage isolated from phage peptide display libraries. J Mol Biol (1994) 4: 639–652.

8. Tal, P. et al. (2016) Cancer therapeutic approach based on conformational stabilization of mutant p53 protein by small peptides. Oncotarget 7:11817–11838.

9. Tal P, Oren M, Palgi O, Rotter V, JEONG K. Development of lead mutant-p53 reactivating peptides towards clinical application [abstract]. Proceedings of the American Association for Cancer Research Annual Meeting 2022; 2022 Apr 8–13. Philadelphia (PA): AACR; Cancer Res 2022;82(12_Suppl):Abstract nr 5288.

10. Ben Abid, S., et al. New Phage Display-Isolated Heptapeptide Recognizing the Regulatory Carboxy-Terminal Domain of Human Tumour Protein p53. Protein J. 2017 Oct;36(5):443–452.

11. Zhu H, Gao H, Ji Y, Zhou Q, Du Z, Tian L, Jiang Y, Yao K, Zhou Z. Targeting p53-MDM2 interaction by small-molecule inhibitors: learning from MDM2 inhibitors in clinical trials. J Hematol Oncol. 2022 Jul 13;15(1):91. doi: 10.1186/s13045-022-01314-3. PMID: 35831864; PMCID: PMC9277894.

12. Isobe, Y. et al. Manumycin polyketides act as molecular glues between UBR7 and P53. Nat Chem Biol. 2020 Nov;16(11):1189–1198. doi: 10.1038/s41589-020-0557-2. Epub 2020 Jun 22. PMID: 32572277; PMCID: PMC7572527.

13. Klein AM, de Queiroz RM, Venkatesh D, Prives C. The roles and regulation of MDM2 and MDMX: it is not just about p53. Genes Dev. 2021 May 1;35(9-10):575–601. doi: 10.1101/gad.347872.120. Epub 2021 Apr 22. PMID: 33888565; PMCID: PMC8091979.

14. Sarafraz-Yazdi E, Mumin S, Cheung D, Fridman D, Lin B, Wong L, Rosal R, Rudolph R, Frenkel M, Thadi A, Morano WF, Bowne WB, Pincus MR, Michl J. PNC-27, a Chimeric p53-Penetratin Peptide Binds to HDM-2 in a p53 Peptide-like Structure, Induces Selective Membrane-Pore Formation and Leads to Cancer Cell Lysis. Biomedicines. 2022 Apr 20;10(5):945. doi: 10.3390/biomedicines10050945. PMID: 35625682; PMCID: PMC9138867.

15. Silva, J.L., Cino, E.A., Soares, I.N., Ferreira, V.F., A P de Oliveira, G. Targeting the Prion-like Aggregation of Mutant p53 to Combat Cancer. Acc Chem Res. 2018; 51:181– 190.

16. Chen, Z., Chen, J., Keshamouni, V.G., Kanapathipillai, M. Polyarginine and its analogues inhibit p53 mutant aggregation and cancer cell proliferation in vitro. Biochemical and Biophysical Research Communications Volume 489, Issue 2, 22 July 2017, Pages 130-134.

17. Wang, G.Z and Fersht, A.R. Multisite aggregation of p53 and implications for drug rescue. Proc Natl Acad Sci U S A. 2017 Mar 28; 114(13): E2634–E2643.

18. Soragni, A. et al. A Designed Inhibitor of p53 Aggregation Rescues p53 Tumor Suppression in Ovarian Carcinomas. Cancer Cell. 2016; 29:90–103.

19. Friedler A, DeDecker BS, Freund SM, Blair C, Rüdiger S, Fersht AR. Structural distortion of p53 by the mutation R249S and its rescue by a designed peptide: implications for “mutant conformation”. J Mol Biol. 2004 Feb 6;336(1):187–96. doi: 10.1016/j.jmb.2003.12.005. PMID: 14741214.

20. Blanden AR, Yu X, Loh SN, Levine AJ, Carpizo DR. Reactivating mutant p53 using small molecules as zinc metallochaperones: awakening a sleeping giant in cancer. Drug Discov Today. 2015 Nov;20(11):1391–7. doi: 10.1016/j.drudis.2015.07.006. Epub 2015 Jul 20. Erratum in: Drug Discov Today. 2016 Oct;21(10):1728. PMID: 26205328; PMCID: PMC4922747.

21. Garufi, A., Ubertini, V., Mancini, F. et al. The beneficial effect of Zinc(II) on low-dose chemotherapeutic sensitivity involves p53 activation in wild-type p53-carrying colorectal cancer cells. J Exp Clin Cancer Res 34, 87 (2015). 10.1186/s13046-015-0206-x

22. Kogan, S., Carpizo, D.R. Zinc Metallochaperones as Mutant p53 Reactivators: A New Paradigm in Cancer Therapeutics. Cancers (Basel). 2018 May 29;10(6):166. doi: 10.3390/cancers10060166. PMID: 29843463; PMCID: PMC6025018.

23. Matochko, W. L., Li S.C, Tang, S.K.Y., and Derda, R. Prospective identification of parasitc sequences in phage display screens. Nucleic Acids Res.2014 Feb, 42(3):1784–98.

24. Braun R, Schönberger N, Vinke S, Lederer F, Kalinowski J, Pollmann K. Application of Next Generation Sequencing (NGS) in Phage Displayed Peptide Selection to Support the Identification of Arsenic-Binding Motifs. Viruses. 2020 Nov 27;12(12):1360. doi: 10.3390/v12121360. PMID: 33261041; PMCID: PMC7759992.

25. Piskacek M, Havelka M, Rezacova M, Knight A. The 9aaTAD Transactivation Domains: From Gal4 to p53. PLoS One. 2016 Sep 12;11(9):e0162842. doi: 10.1371/journal.pone.0162842. PMID: 27618436; PMCID: PMC5019370.

26. Barbas, C.F., Burton, D.R., Scott, J.K., Silverman, G.J. (2001) Phage display: a laboratory manual. Cold Spring Harbor Laboratory Press, New York.

27. Rowell, J.P., Simpson, K.L., Stott, K., Watson, M., Thomas, J.O. HMGB1-facilitated p53 DNA binding occurs via HMG-Box/p53 transactivation domain interaction, regulated by the acidic tail. Structure. 2012 Dec 5;20(12):2014–24. doi: 10.1016/j.str.2012.09.004. Epub 2012 Oct 11. PMID: 23063560.

28. Petty, T.J. et al. An induced fit mechanism regulates p53 DNA binding kinetics to confer sequence specificity. EMBO J. 2011 Jun 1;30(11):2167–76. doi: 10.1038/emboj.2011.127. Epub 2011 Apr 26. PMID: 21522129; PMCID: PMC3117648.

29. Trott, O., Olson, A.J. (2010). AutoDock Vina: improving the speed and accuracy of docking with a new scoring function, efficient optimization, and multithreading. J. computat chem 31(2), 455–461.

30. Sloth AB, Bakhshinejad B, Jensen M, Stavnsbjerg C, Liisberg MB, Rossing M, Kjaer A. Analysis of Compositional Bias in a Commercial Phage Display Peptide Library by Next-Generation Sequencing. Viruses. 2022 Oct 29;14(11):2402. doi: 10.3390/v14112402. PMID: 36366500; PMCID: PMC9697088.

31. Luck K, Travé G (2011) Phage display can select overhydrophobic sequences that may impair prediction of natural domain–peptide interactions. Bioinf Discov 27:899–902.

32. Ma L, Green SI, Trautner BW, Ramig RF, Maresso AW. Metals Enhance the Killing of Bacteria by Bacteriophage in Human Blood. Sci Rep. 2018 Feb 2;8(1):2326. doi: 10.1038/s41598-018-20698-2. PMID: 29396496; PMCID: PMC5797145.

33. Nagata, T. et al. (2014) Structural Basis for Inhibition of the MDM2:p53 Interaction by an Optimized MDM2-Binding Peptide Selected with mRNA Display. PLoS ONE 9(10): e109163. 10.1371/journal.pone.0109163.

34. Blanden, A.R. et al. Zinc shapes the folding landscape of p53 and establishes a pathway for reactivating structurally diverse cancer mutants. Elife. 2020 Dec 2;9:e61487. doi: 10.7554/eLife.61487. PMID: 33263541; PMCID: PMC7728444.

35. Ha, J.H., Prela, O., Carpizo, D.R., Loh, S.N. p53 and Zinc: A Malleable Relationship. Front Mol Biosci. 2022 Apr 13;9:895887. doi: 10.3389/fmolb.2022.895887. PMID: 35495631; PMCID: PMC9043292.

36. Bochkareva E, Kaustov L, Ayed A, Yi G-S, Lu Y, Pineda-Lucena A, Liao JCC, Okorokov AL, Milner J, Arrowsmith CH, Bochkarev A. 2005. Single-stranded DNA mimicry in the p53 transactivation domain interaction with replication protein A. Proc Natl Acad Sci USA 102:15412–15417.

37. Liu X, Chen J. Modulation of p53 Transactivation Domain Conformations by Ligand Binding and Cancer-Associated Mutations. Pac Symp Biocomput. 2020;25:195–206. PMID: 31797597; PMCID: PMC6934143.

