## Supplementary material for "Phage libraries screening on P53 : yield improvement by zinc and a new parasites-integrating analysis and rationale": SD

| peptide | sequence | E (2LY4B) | E (3Q01) |
|---|---|---|---|
| 7.1 | HTWLRSA | -5,4 | -5,8 |
| 7.2 | LHNSLPA | -4,7 | -4,7 |
| 7.3 | NPNSAQG | -3,9 | -5,7 |
| 7.4 | ATHQTLR | -3,7 | -5,6 |
| 7Z1 | WSWPRFL | -4,9 | -5,9 |
| 7Z2 | MQAPSPM | -4,1 | -5,8 |
| 7Z3 | AAAFTQS | -4,8 | -5,7 |
| 7Z4 | GTEPPAM | -4,3 | -5,5 |
| 12.1 | NHMNQISFPSRP | -4,3 | -5,6 |
| 12.2 | ARSPCQVQSRTS | -4,1 | -4,1 |
| 12.3 | NNLAFYHTFISP | -4,6 | -6,8 |
| 12.4 | APSPFQVQSRTS | -5,9 | -6,4 |
| 12.5 | NYPSSSVPHAPQ | -4,5 | -6,9 |
| 12.6 | YSTHDNARPWLL | -5 | -5,3 |
| 12Z1 | SHVPLARWSVIT | -4,8 | -6 |
| 12Z2 | HDHLIPFYWADL | -5,2 | -6 |
| 12Z3 | STLVFPAHTRDY | -4,2 | -4,9 |
| 12Z4 | TYLLPHSYPWYG | -4,6 | -6,8 |
| 12Z5 | TATLDMPLSLPS | -3,9 | -6,2 |
| 12Z6 | WMDSYMSQHDWP | -4,3 | -5 |
| PD.1 | GANMKYA | -3,7 | -5,4 |
| PD.2 | GLTATNM | -4,7 | -5 |
| PD.3 | GFTATNM | -5,1 | -5,8 |
| PD.4 | NDAEMPT | -2,8 | -5,6 |
| PD.5 | ETTHARA | -3,9 | -6,4 |
| PD.6 | GLDCYKQ | -3,7 | -5,9 |
| PD.7 | STQARTP | -4,9 | -6,1 |
| SR12.1 | HLAQTASPPAAP | -4,9 | -6,3 |
| SR12.2 | APLYSPSHLATS | -5,7 | -5,9 |

a

| Pep | Sequence | E (2LY4B) |
|---|---|---|
| R0 | VGVRIPL | -4,2 |
| R1 | NGVEIPP | -5 |
| R3 | PFNEPHP | -5,4 |
| R4 | PINEPHP | -5,5 |
| R6 | SFNEPHP | -4,8 |
| R7 | SINEPHP | -4,7 |

b

| Pep | Sequence | E (3Q01) |
|---|---|---|
| R0 | VGVRIPL | -6,2 |
| R1 | NGVEIPP | -6,8 |
| R3 | PFNEPHP | -6,9 |
| R4 | PINEPHP | -6,7 |
| R2 | PFNEPHL | -7 |
| R5 | PKNEPHP | -7,1 |
| R8 | VGVGIPP | -5,9 |
| R9 | PGVGIPL | -5,9 |
| R10 | IRVGIPL | -6,2 |
| R11 | LFNERHP | -6 |
| R12 | PINEPHL | -6,7 |
| R13 | AFNEPHP | -5,7 |
| R14 | AINEPHP | -5,9 |
| R15 | AINEPHL | -6 |
| R16 | AFHEPHP | -7,4 |
| R17 | AIHEPHP | -5,9 |

c

**Table SD1:** Energy of docking. a) all peptides except those of R set. b) R set representatives for 2LY4B docking. And c) R set representatives for 3Q01 docking.

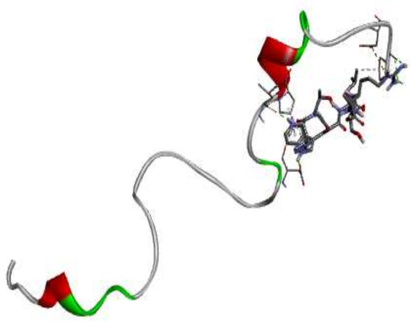

7.1: HTWLRSA

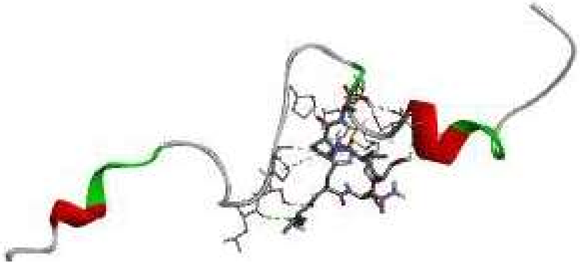

7.2 : LHNSLPA

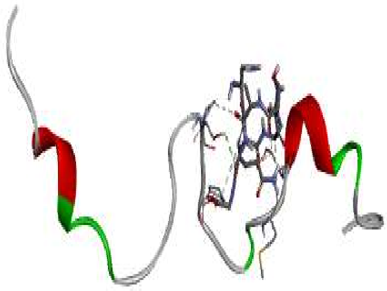

7.3: NPNSAQG

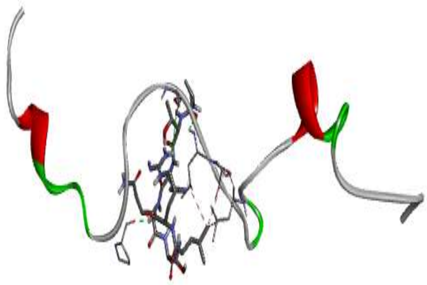

7.4 : ATHQTLR

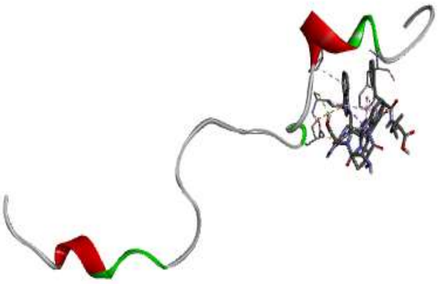

7Z1 : WSWPRFL

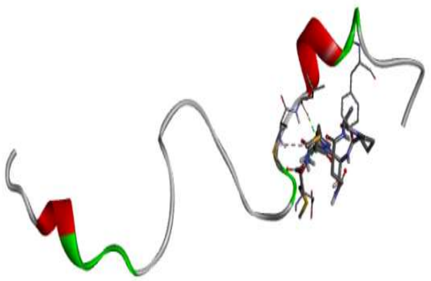

7Z2 : MQAPSPM

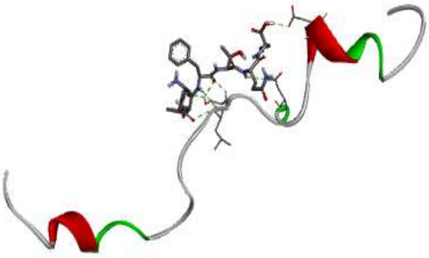

7Z3 : AAAFTQS

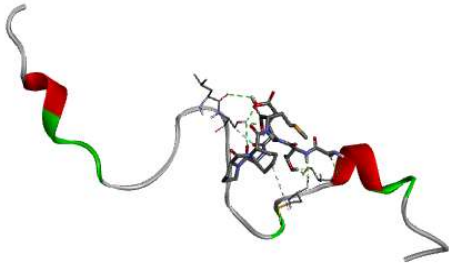

7Z4 : GTEPPAM

**Figure 1.1.SD : Docking structures of 7-mer set with 2LY4.B.**

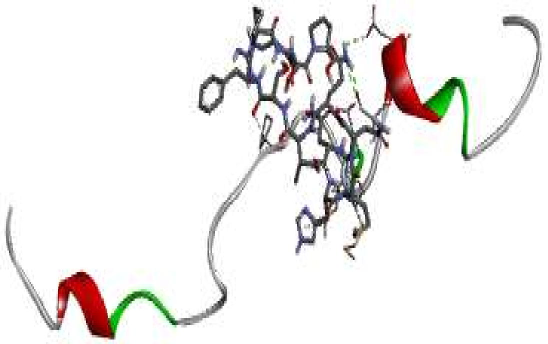

12.1 : NHMNQISFPSRP

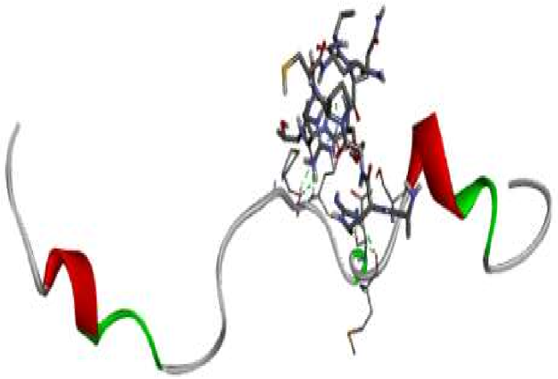

12.2: ARSPCQVQSRTS

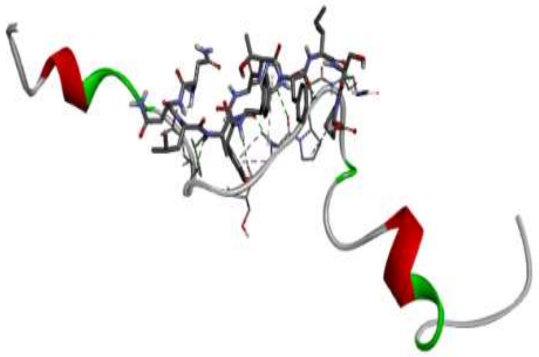

12.3: NNLAFYHTFISP

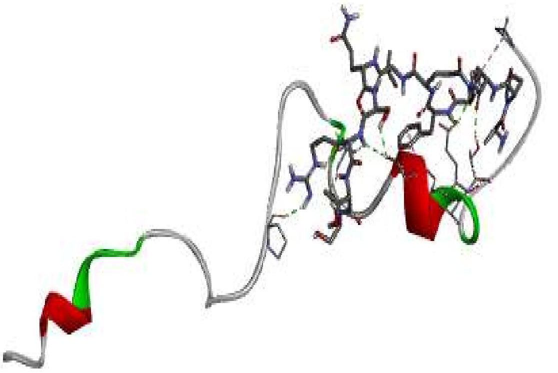

12.4 : APSPFQVQSRTS

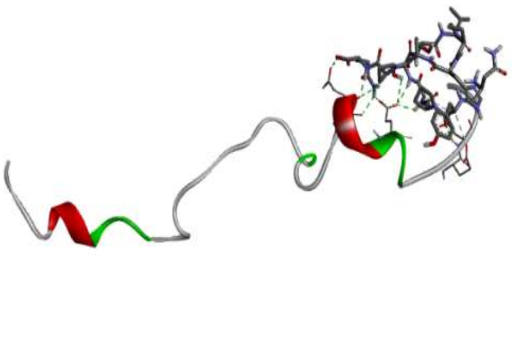

12.5: NYPSSSVPHAPQ

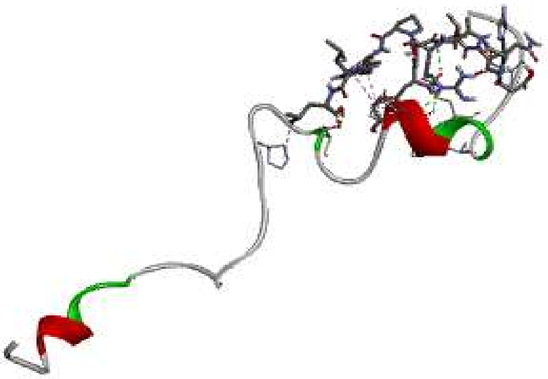

12.6: YSTHDNARPWLL

**Figure 1.2.SD: Docking structures of 12-mer “*Non Zinc*” set with 2LY4.B.**

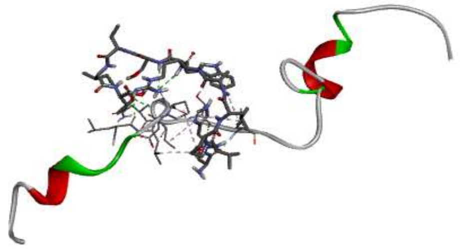

12z1:SHVPLARWSVIT

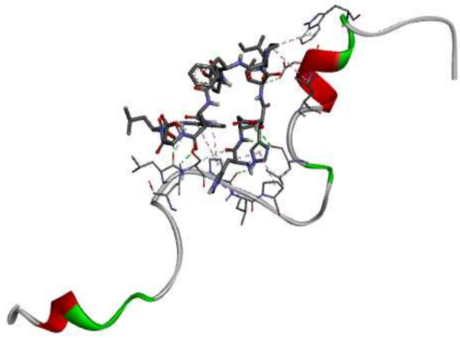

12z2: HDHLIPFYWADL

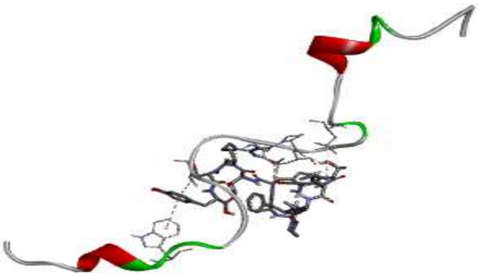

12z3:STLVFPAHTRDY

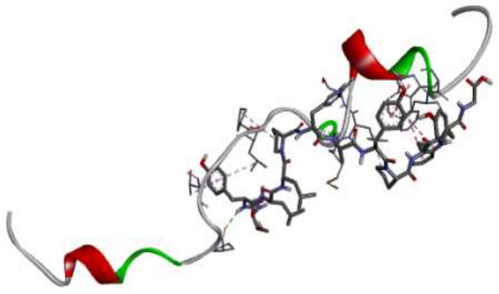

12z4:TYLLPHSYPWYG

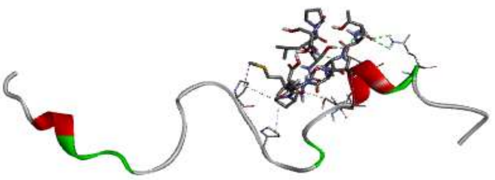

12z5: TATLDMPLSLPS

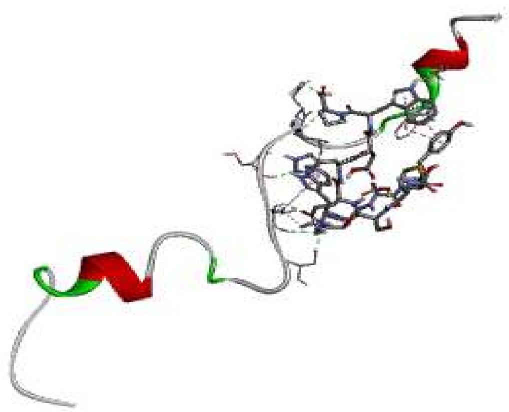

12z6:WMDSYMSQHDWP

**Figure 1.3.SD: Docking structures of 12-mer *“With Zinc”* set with 2LY4.B.**

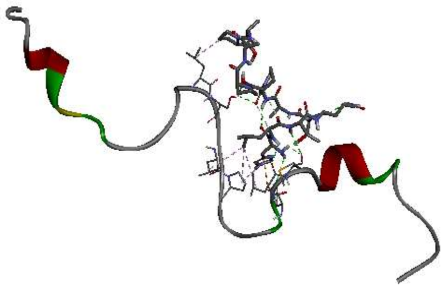

SR12.1 : HLAQTASPPAAP

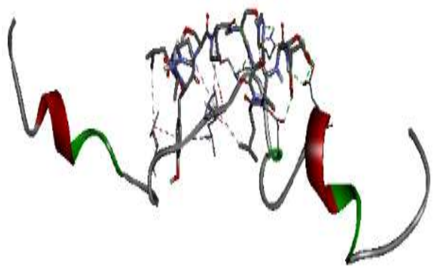

SR12.2 : APLYSPSHLATS

**a. SR50 peptides.**

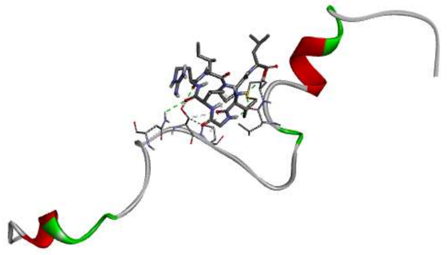

R0 : VGVRIPL

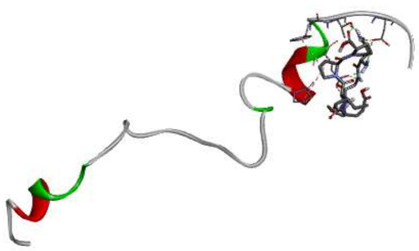

R1 : NGVEIPP

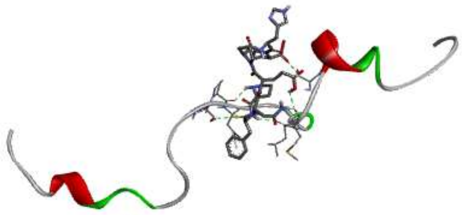

R2 : PFNEPHP

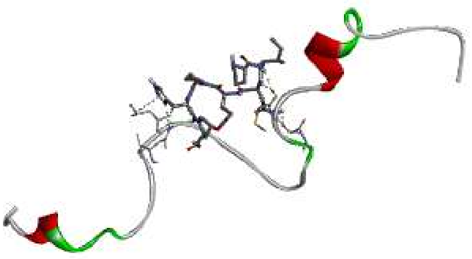

R3 : PINEPHP

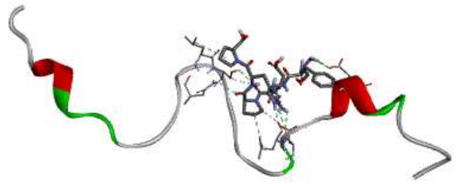

R6 : SFNEPHP

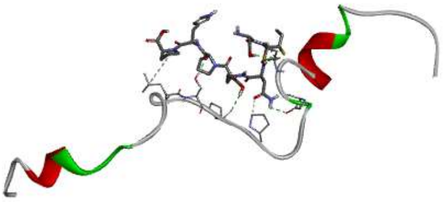

R7 : SINEPHP

**b. Redundant (R) peptides.**

**Figure 1.5.SD: Docking structures of SR50 and Redundant (R) sets with 2LY4.B.**

a. SR50 peptides. b. Representatives of Redundant (R) peptides.

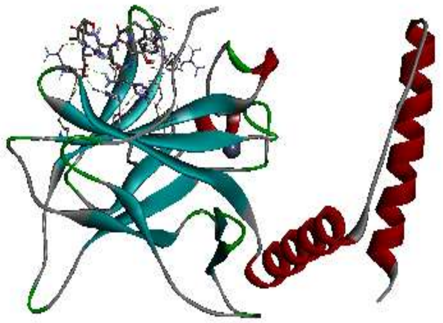

7.1: HTWLRSA

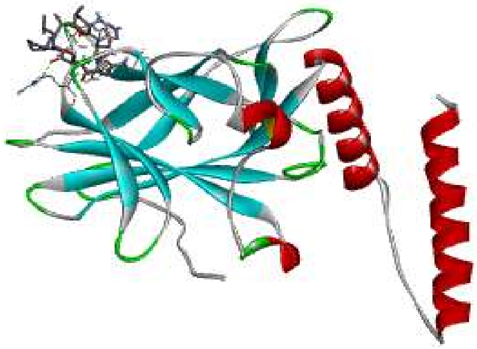

7.2: LHNSLPA

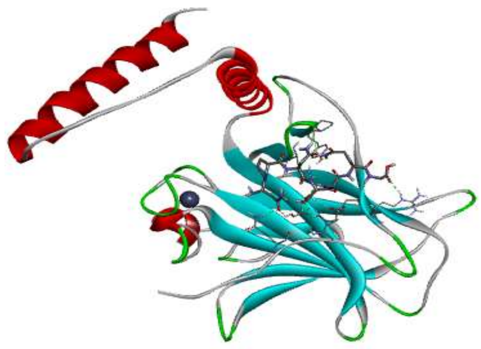

7.3: NPNSAQG

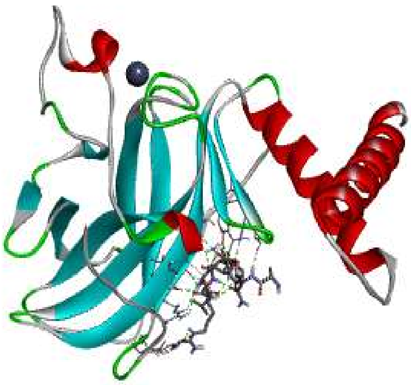

7.4: ATHQTLR

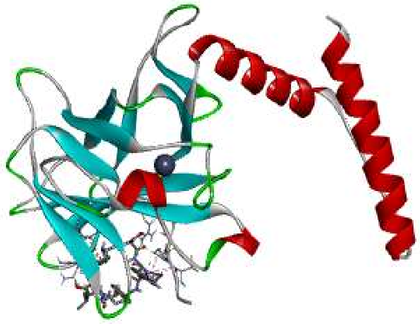

7Z1:WSWPRFL

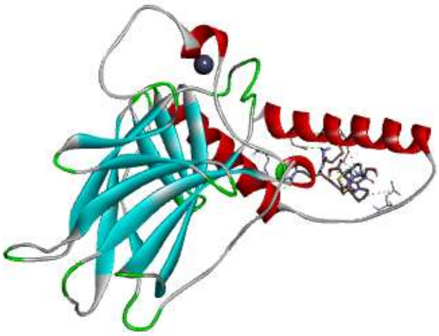

7Z2: MQAPSPM

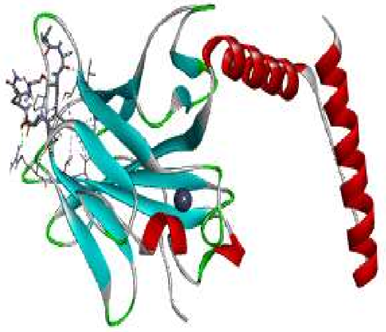

7Z3:AAAFTQS

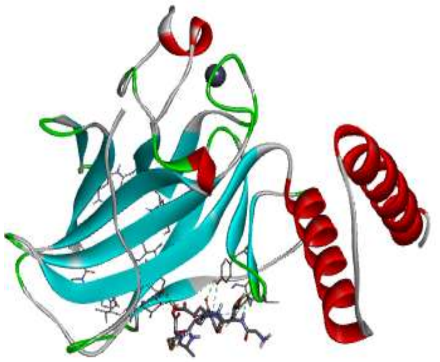

7Z4:GTEPPAM

**Figure 2.1.SD : Docking structures of *7-mer* set with 3Q01 (ribbon).**

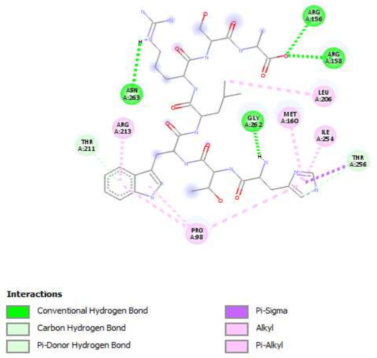


7.1: HTWLRSA

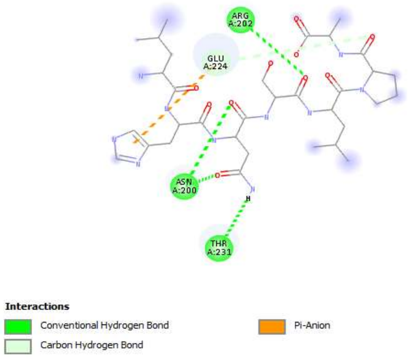


7.2: LHNSLPA

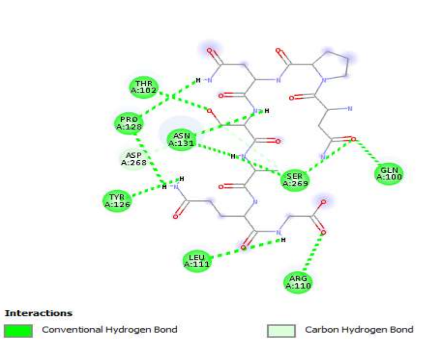


7.3: NPNSAQG

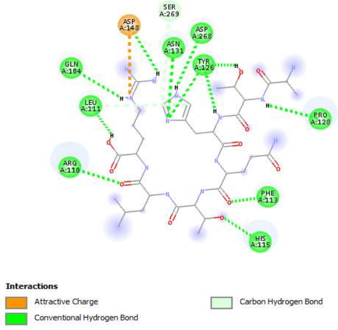


7.4: ATHQTLR

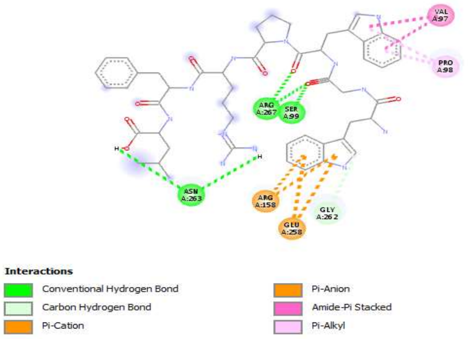


7Z1:WSWPRFL

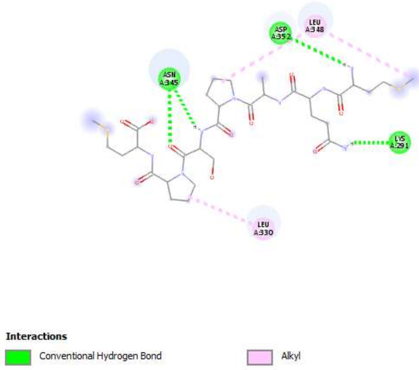


7Z2: MQAPSPM

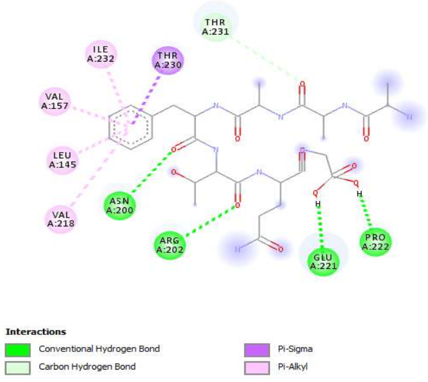


7Z3:AAAFTQS

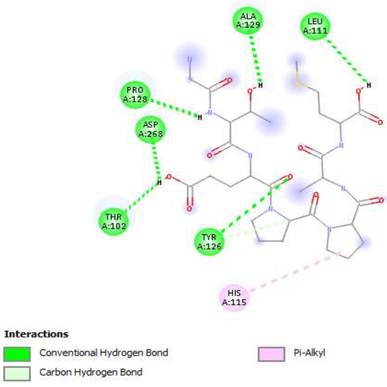


7Z4: GTEPPAM

**Figure 2.2.SD : Docking structures of *7-mer* set with 3Q01 (interactions).**

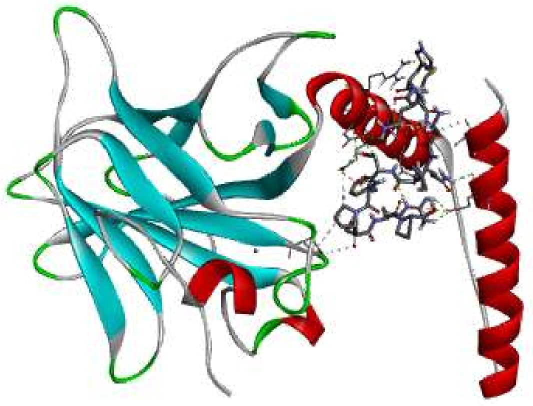

12.1 : NHMNQISFPSRP

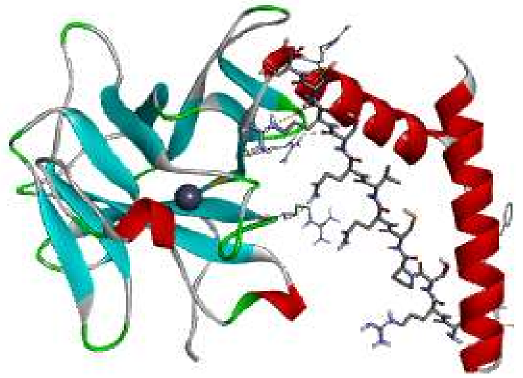

12.2 : ARSPCQVQSRTS

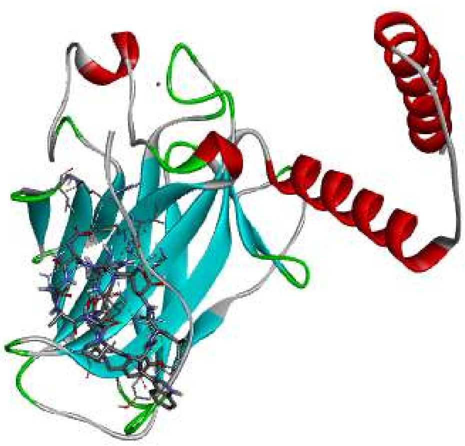

12.6:YSTHDNARPWLL

**Figure 2.3.SD : Docking structures of 12-mer *non zinc* set with 3Q01 (ribbon).**

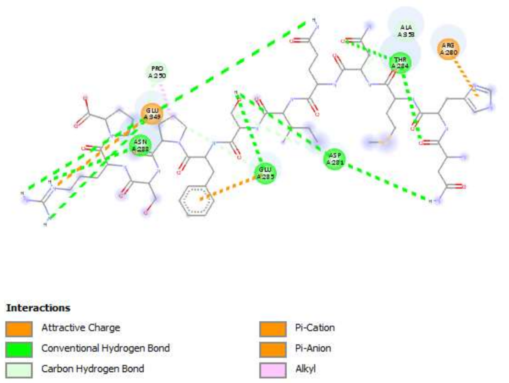


12.1: NHMNQISFPSRP

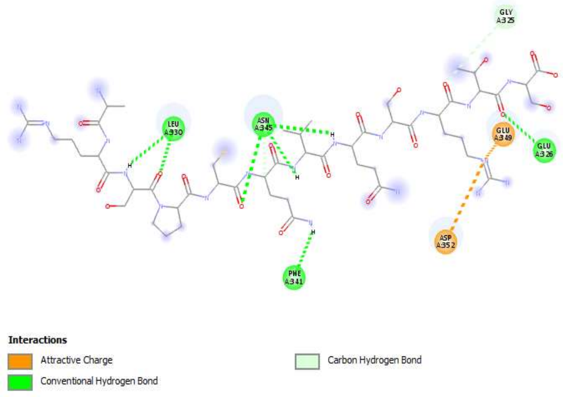


12.2: ARSPCQVQSRTS

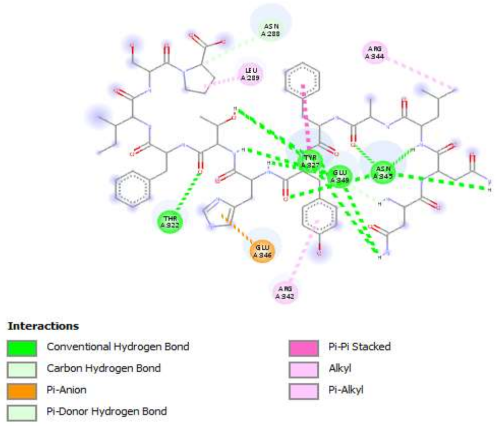


12.3: NNLAFYHTFISP

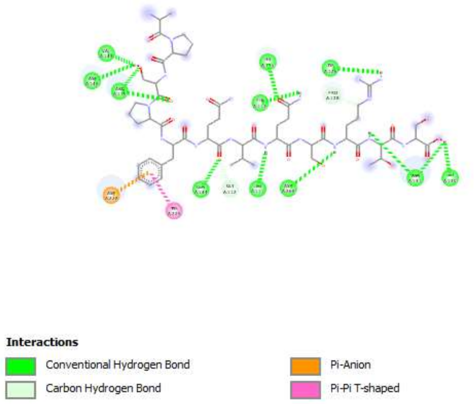


12.4: APSPFQVQSRTS

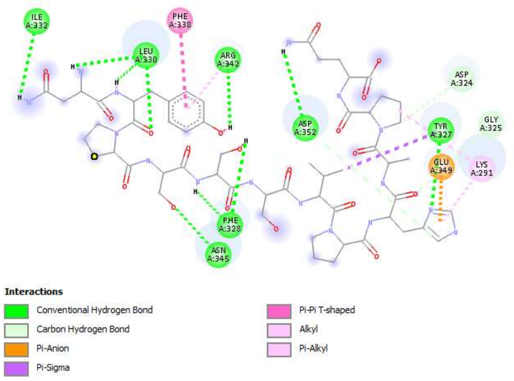


12.5: NYPSSSVPHAPQ

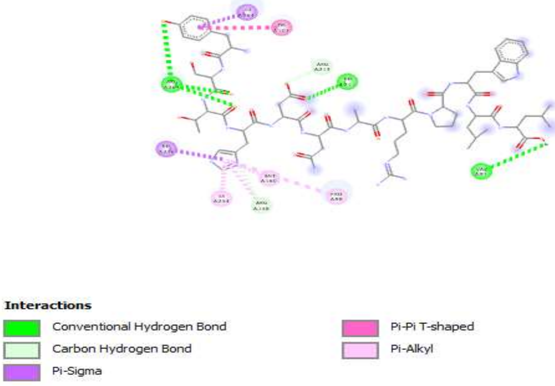


12.6:YSTHDNARPWLL

**Figure 2.4.SD: Docking structures of 12-mer *non zinc* set with 3Q01 (interactions).**

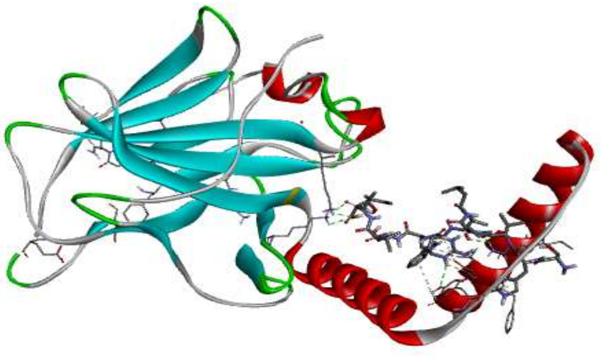

12Z1:SHVPLARWSVIT

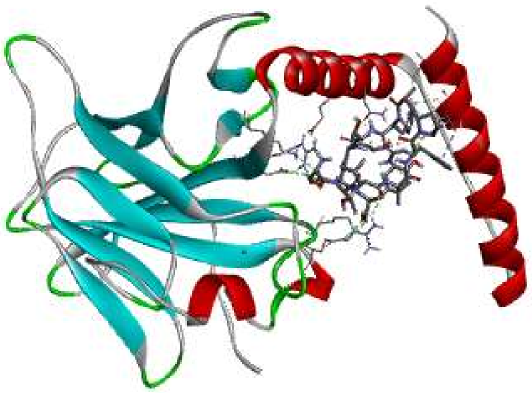

12Z3:STLVFPAHTRDY

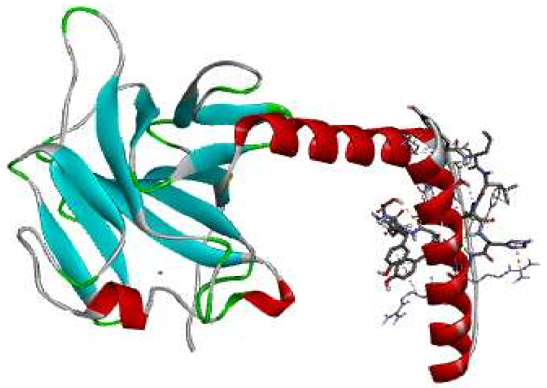

12Z4:TYLLPHSYPWYG

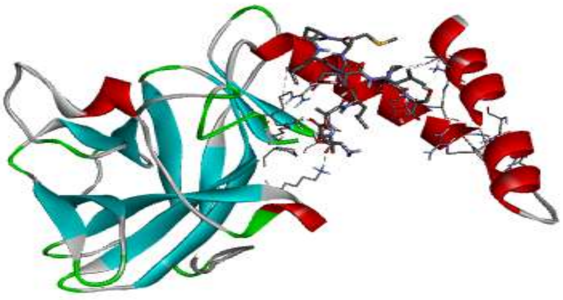

12Z5:TATLDMPLSLPS

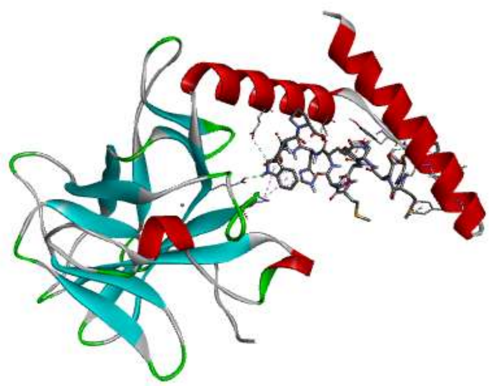

12Z6:WMDSYMSQHDWP

**Figure 2.5.SD: Docking structures of 12-mer *with zinc* set with 3Q01 (ribbon).**

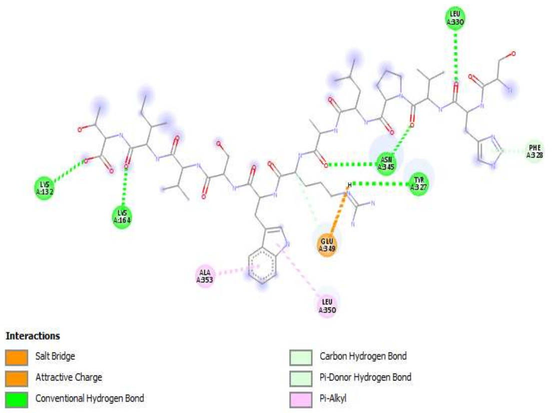


12Z1: SHVPLARWSVIT

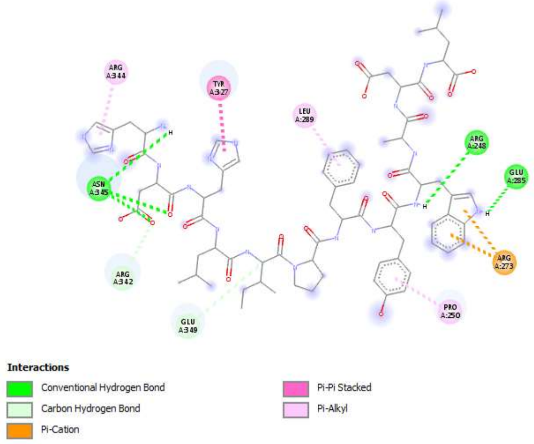


12Z2: HDHLIPFYWADL

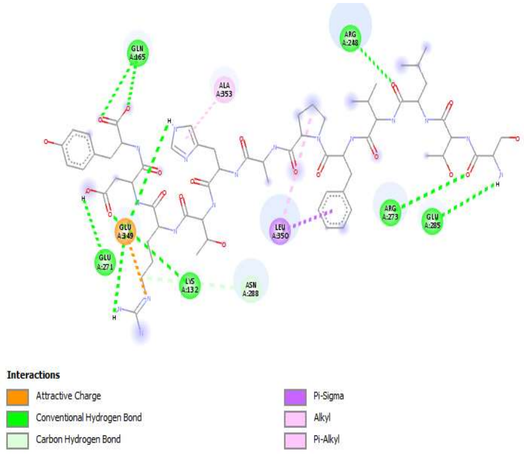


12Z3: STLVFPAHTRDY

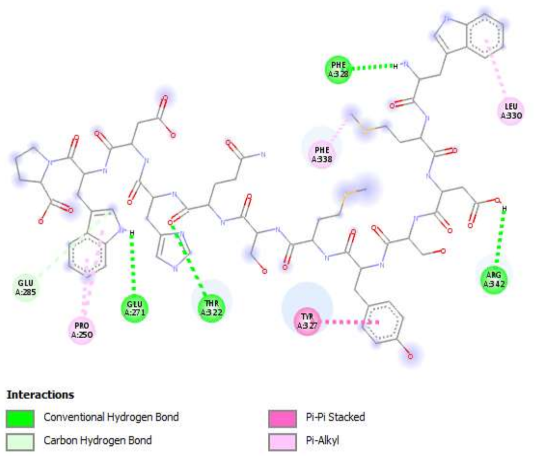


12Z6: WMDSYMSQHDWP

**Figure 2.6.SD: Docking structures of 12-mer *with zinc* set with 3Q01 (interactions).**

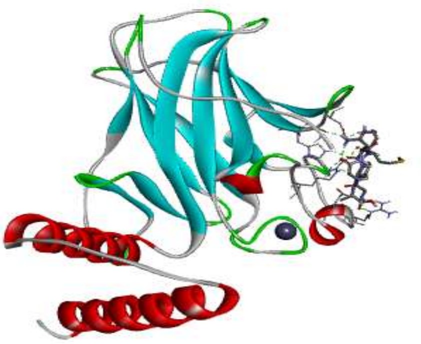

PD1 : GANMKYA

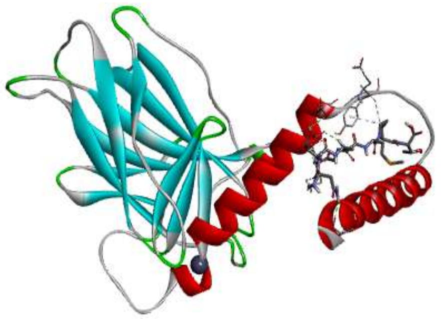

PD.4 : NDAEMPT

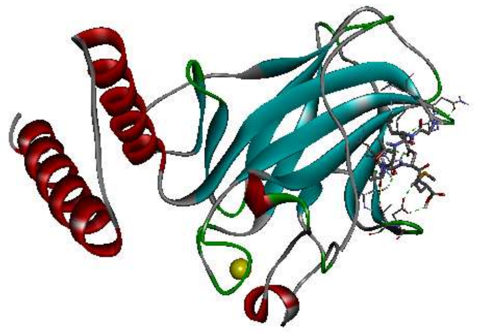

PD2: GLTATNM

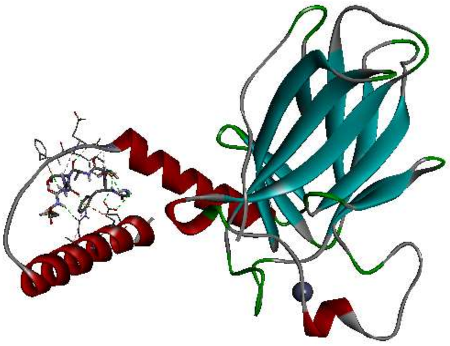

PD3: GFTATNM

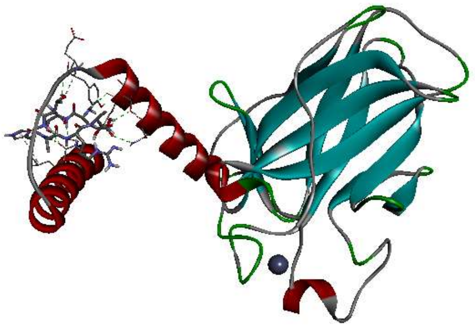

PD5: ETTHARA

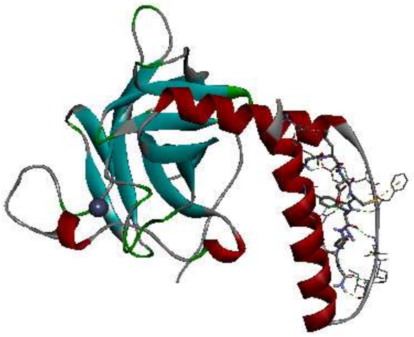

PD6: GLDCYKQ

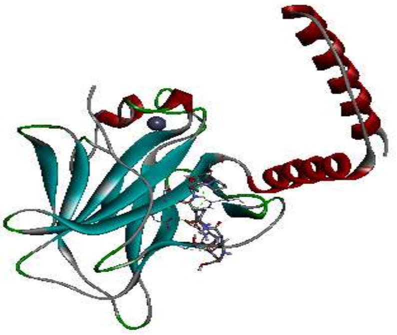

PD7: STQARTP

**Figure 2.7.SD: Docking structures of PD74 set (control) with 3Q01 (ribbon).**

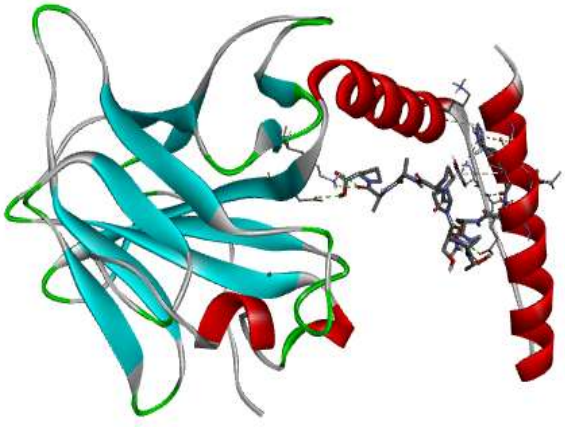

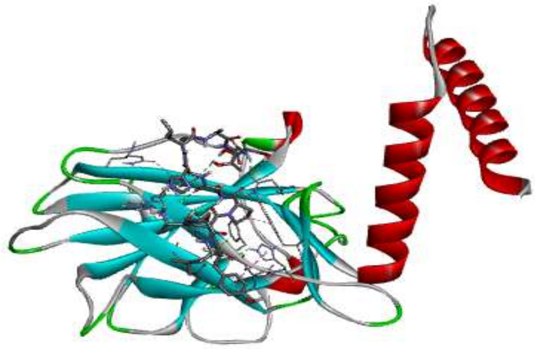

SR12.1: HLAQTASPPAAP SR12.2:APLYSPSHLATS

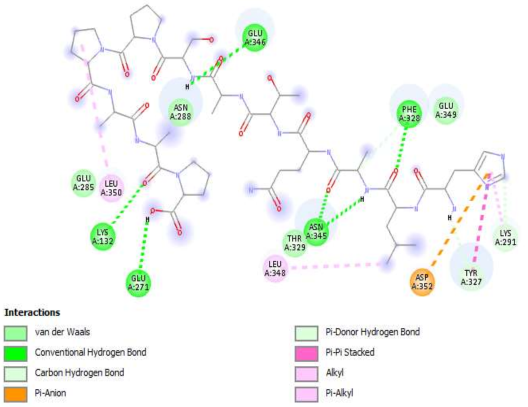


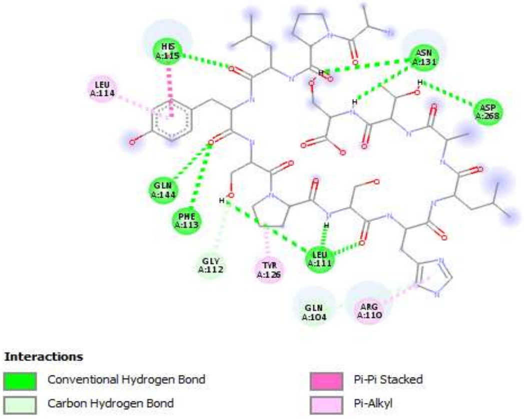


SR12.1: HLAQTASPPAAP SR12.2:APLYSPSHLATS

**Figure 2.8.SD: Docking structures of SR50 set with 3Q01.**

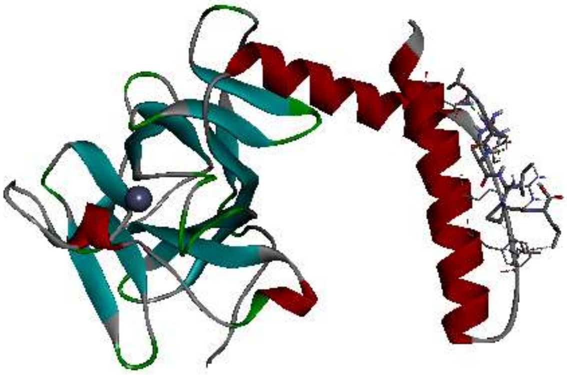

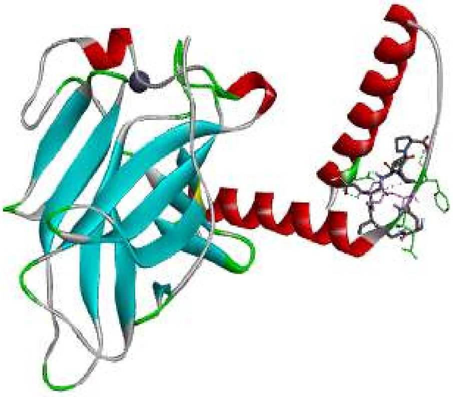

R0: VGVRIPL

R1: NGVEIPP

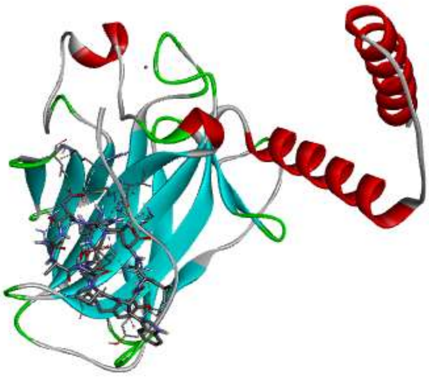

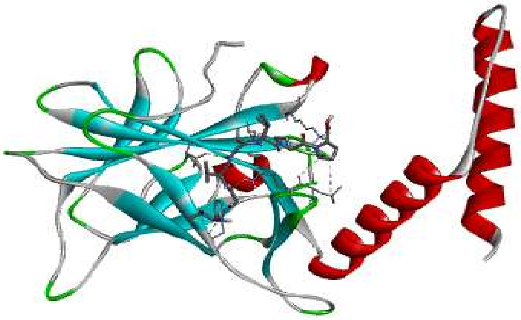

R8:VGVGIPP

R9: PGVGIPL

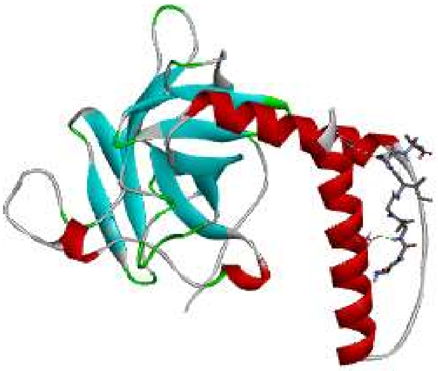

R10: IRVGIPL

**Figure 2.9.SD : Docking structures of Redundant set (R) Motif 1 with 3Q01 (ribbon).**

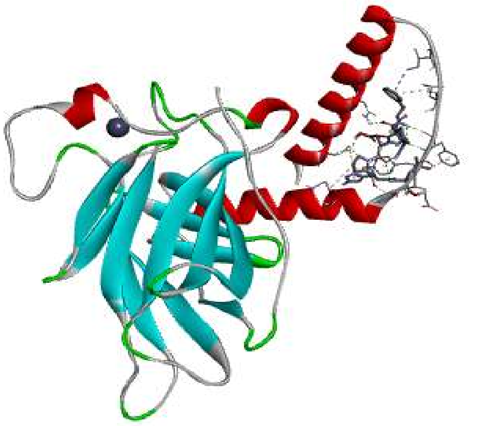

R2: PFNEPHL

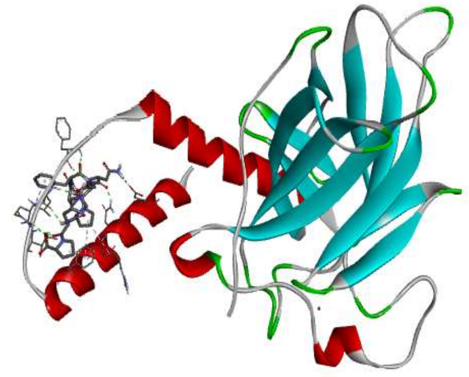

R3: PFNEPHP

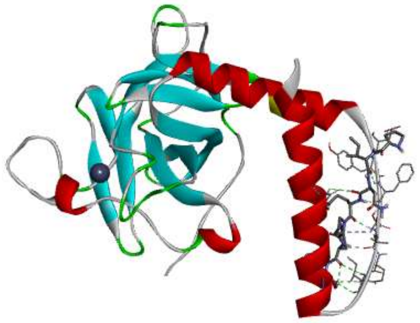

R4: PINEPHP

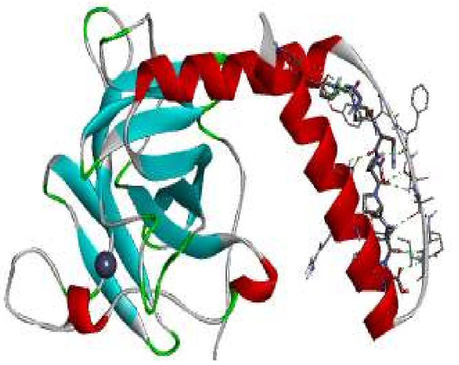

R5: PKNEPHP

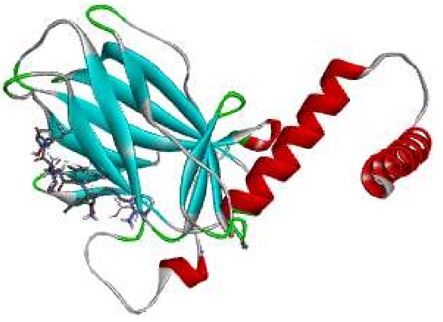

R11: LFNERHP

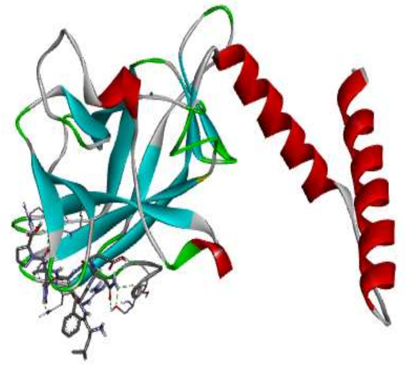

R12: PINEPHL

**Figure 2.10.SD: Docking structures of Redundant set (R) Motif 2 with 3Q01 (ribbon).**

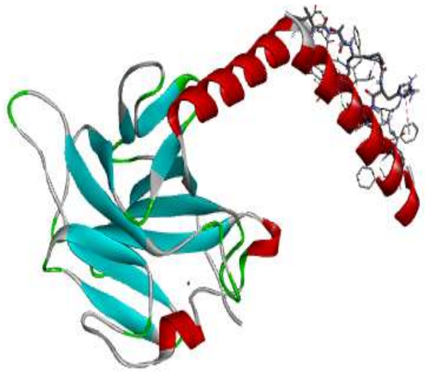

R13 : AFNEPHP

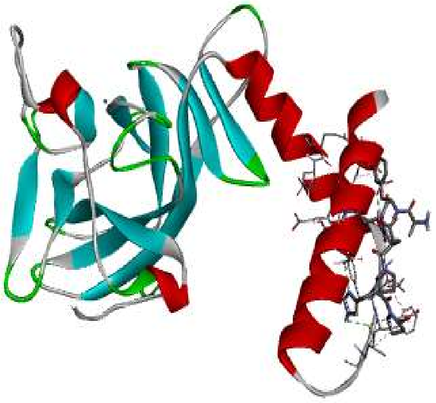

R14:AINEPHP

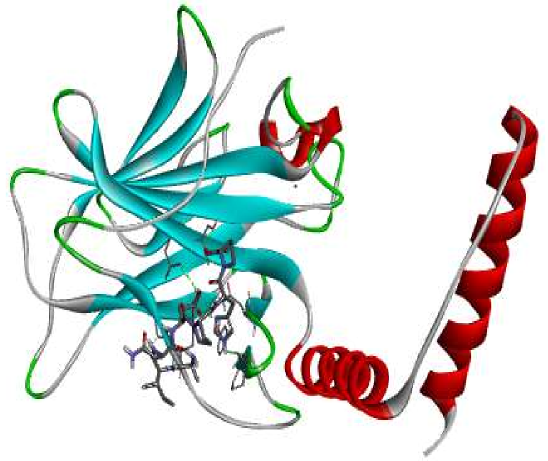

R15: AINEPHL

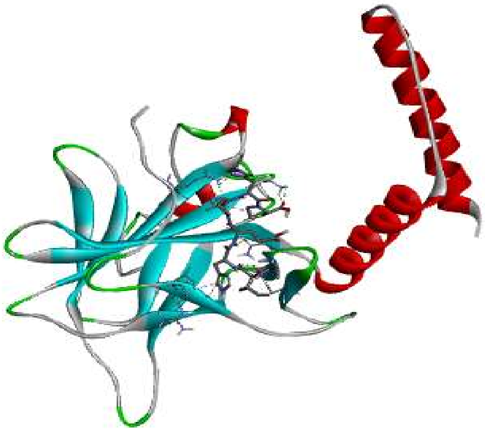

R16:AFHEPHP

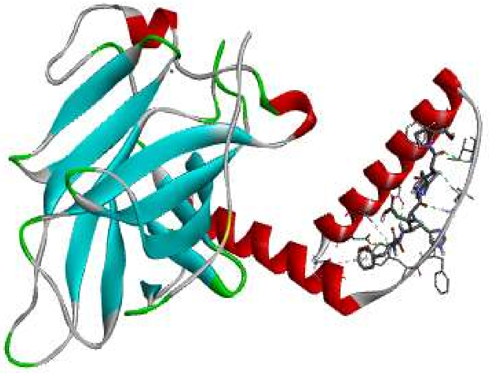

R17:AIHEPHP

**Figure 2.11.SD: Docking structures of Redundant set (R) Motif 2 with 3Q01 (ribbon) (continuation).**

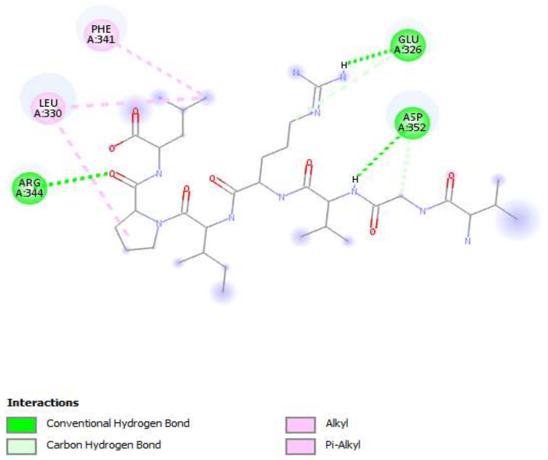


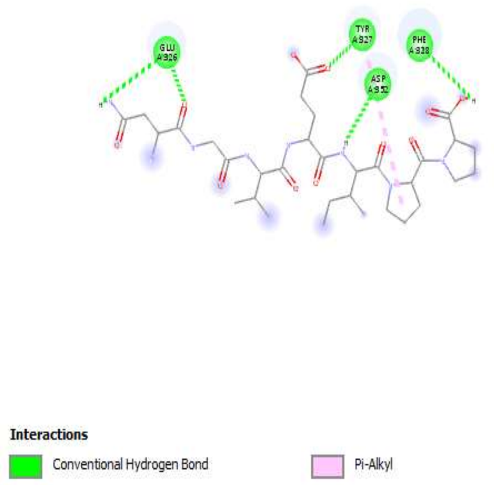


R0: VGVRIPL

R1: NGVEIPP

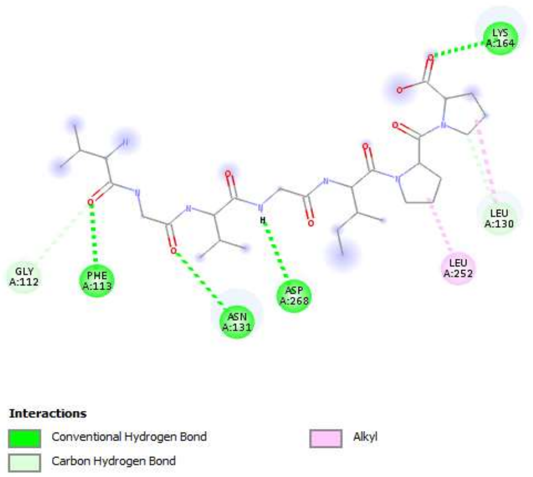


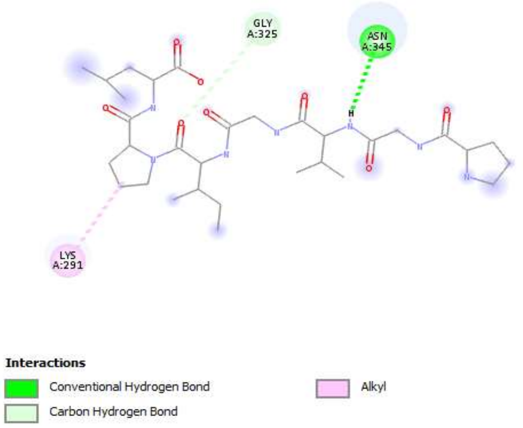


R8:VGVGIPP

R9: PGVGIPL

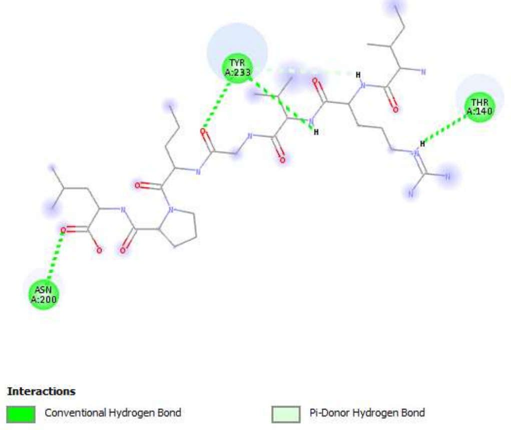


R10: IRVGIPL

**Figure 2.12.SD : Docking structures of Redundant set (R) Motif 1 with 3Q01 (interactions).**

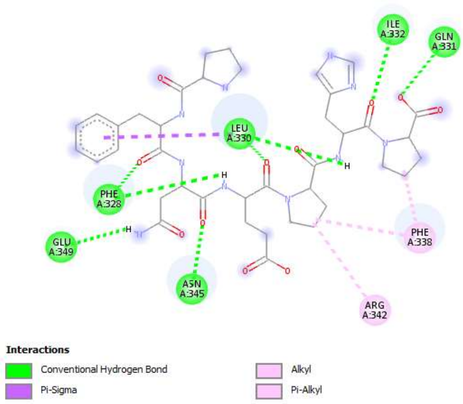


R3: PFNEPHP

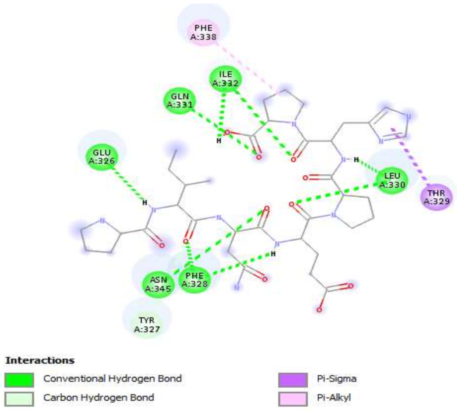


R4: PINEPHP

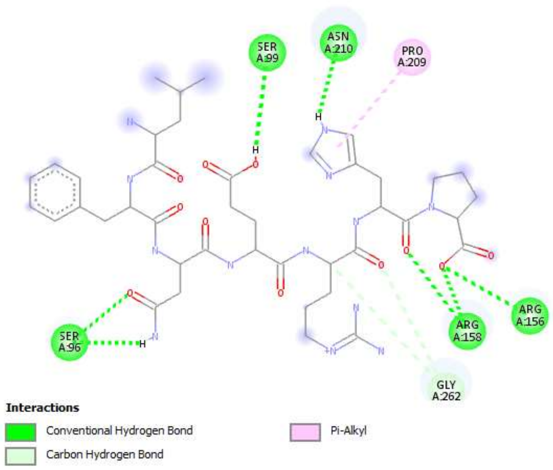


R11: LFNERHP

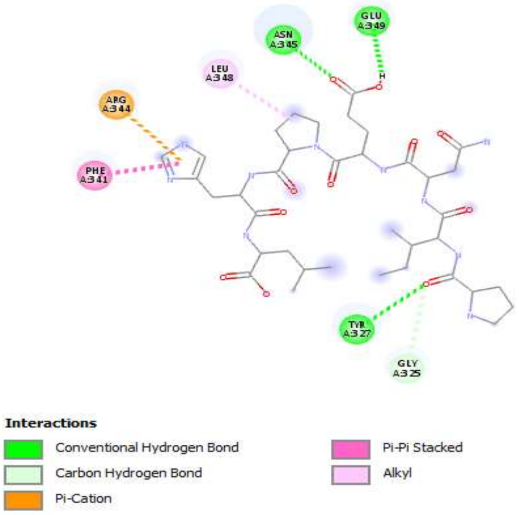


R12: PINEPHL

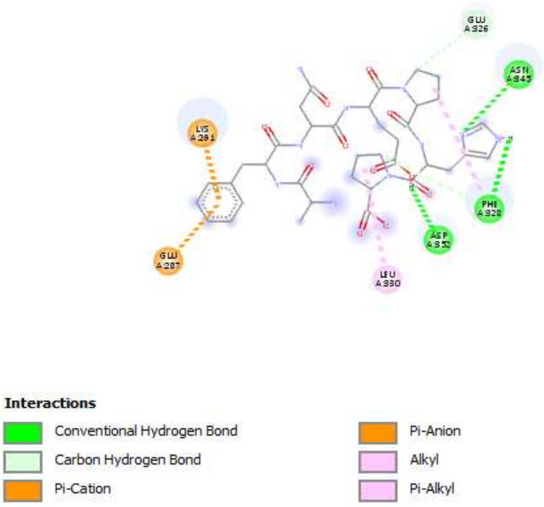


R13 : AFNEPHP

**Figure 2.13.SD: Docking structures of Redundant set (R) Motif 2 with 3Q01 (interactions).**

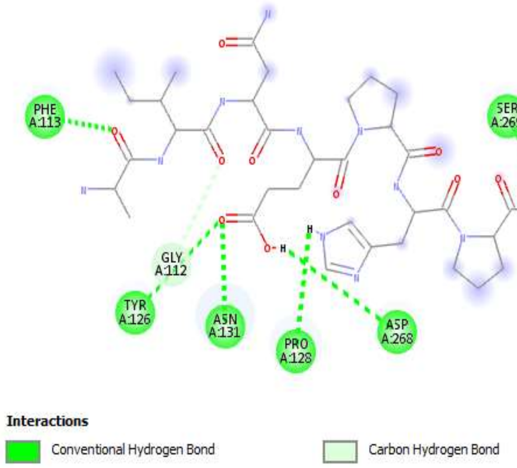


R14:AINEPHP

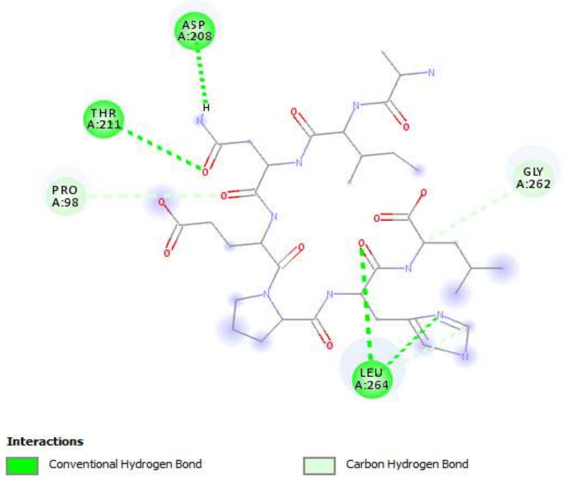


R15: AINEPHL

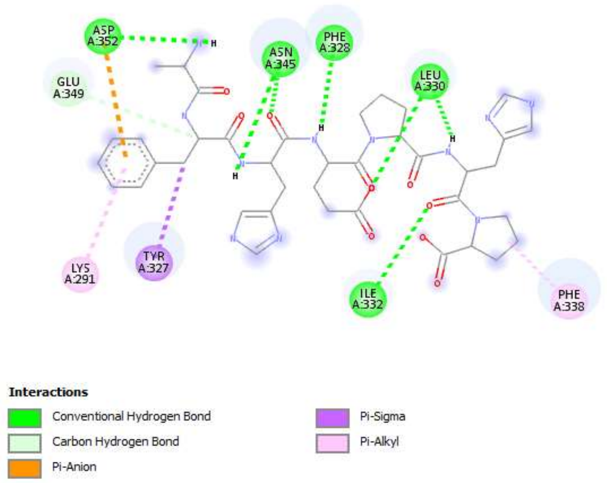


R16:AFHEPHP

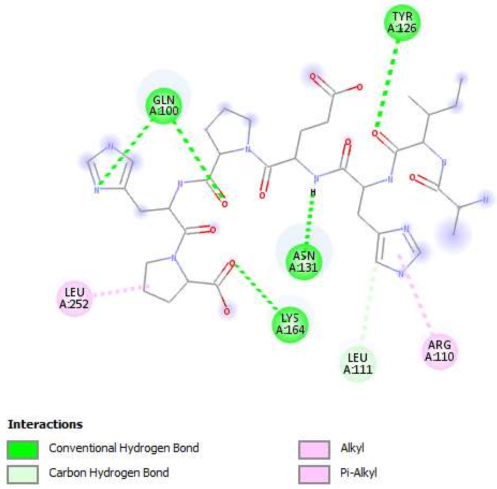


R17:AIHEPHP

**Figure 2.14.SD: Docking structures of Redundant set (R) Motif 2 with 3Q01 (interactions) (continuation).**

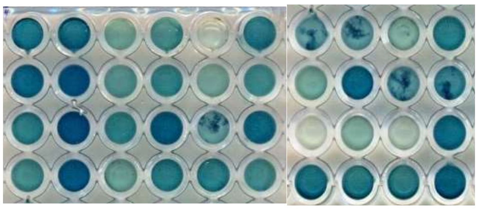

| C | P | C | P | C | P | C | P | C | P |
|---|---|---|---|---|---|---|---|---|---|
| 7.4 | 7.4 | 7Z3 | 7Z3 | 12Z4 | 12Z4 | 12Z1 | 12Z1 | 12.1 | 12.1 |
| PD1 | PD1 | 7Z1 | 7Z1 | 12.6 | 12.6 | 7.2 | 7.2 | 12Z2 | 12Z2 |
| 12Z5 | 12Z5 | 12.3 | 12.3 | 7Z4 | 7Z4 | 7.3 | 7.3 | 12.4 | 12.4 |
| S12.2 | S12.2 | PD7 | PD7 | 12Z3 | 12Z3 | 12.5 | 12.5 | 7.1 | 7.1 |

**a**

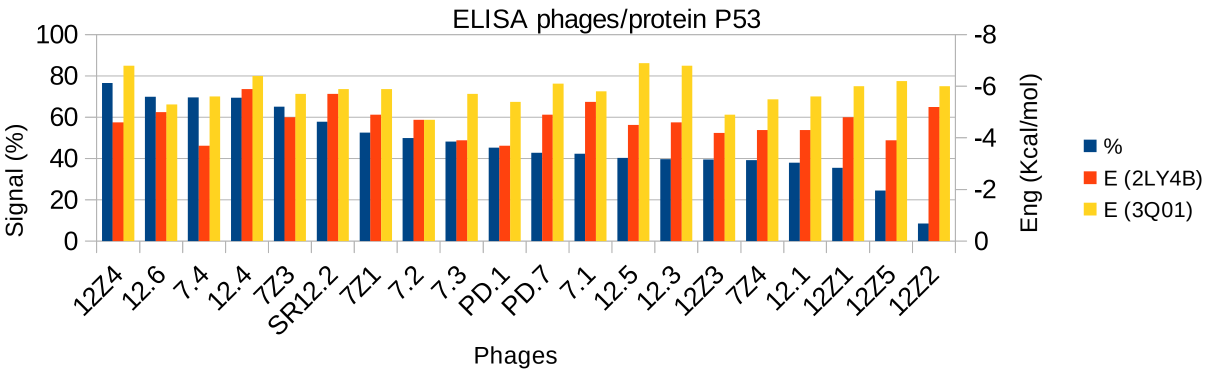


**b**

**Fig.3.SD. ELISA of phages on protein P53.**

**a)** Photo of 96-well plate after revelation and its lay-out aside. **b)** Histogram representing the responses of different tested phage clones. Wells are coated with protein P53. Phages are added separately. Following incubation with phages, anti-M13 antibody-HRP (Cytiva Cat# 27942101, RRID:AB_2616587) is added and response is revealed with HRP substrate (ABTS). Phage clones from phage display experiment against p53-derived peptides are respectively PD1 and PD7 for PD74 (12-61) and SR12.2 for SR50 (241-291).The 96-well plate columns were alternatively coated by protein P53 target (P) or only blocking buffer (C for Control). Each phage is essayed simultaneously on control and protein wells. A percentage of relative signals calculated as: [100-C/P*100]. Clones are ranked based on this percentage value representing the binding force. At secondary y axis: energies of docking on both 2LY4B and 3Q01 structures. This ELISA is Representative.
